# A High-Throughput Microfluidic Quantitative PCR Platform for the Simultaneous Quantification of Pathogens, Fecal Indicator Bacteria, and Microbial Source Tracking Markers

**DOI:** 10.1101/2023.02.25.529995

**Authors:** Elizabeth R. Hill, Chan Lan Chun, Kerry Hamilton, Satoshi Ishii

## Abstract

Contamination of water with bacterial, viral, and protozoan pathogens can cause human diseases. Both humans and non-humans can release these pathogens through their feces. To identify the sources of fecal contamination in the water environment, microbial source tracking (MST) approaches have been developed; however, the relationship between MST markers and pathogens is still not well understood most likely due to the lack of comprehensive datasets of pathogens and MST marker concentrations. In this study, we developed a novel microfluidic quantitative PCR (MFQPCR) platform for the simultaneous quantification of MST markers, fecal indicator bacteria (FIB), and bacterial, viral, and protozoan pathogens in many samples. A total of 80 previously validated TaqMan probe assays were applied on the MFQPCR chips, including those for two FIB, 22 bacterial pathogens, 11 viral pathogens, five protozoan pathogens, 37 MST markers for various host species, and three process controls. Specific and sensitive detection was verified for most assays on the MFQPCR platform. The MFQPCR chip was applied to analyze pathogen removal rates during the wastewater treatment processes. In addition, multiple host-specific MST markers, FIB, and pathogens were successfully quantified in human and avian-impacted surface waters. While the genes for pathogens were relatively infrequently detected, positive correlations were observed between some potential pathogens such as *Clostridium perfringens* and *Mycobacterium* spp., and human MST markers. The MFQPCR chips developed in this study, therefore, can provide useful information to monitor and improve water quality.

## 1. Introduction

Waters contaminated with bacterial, viral, and protozoan pathogens pose an increased risk of infections (Harwood et al., 2014). Currently, fecal indicator bacteria (FIB) such as *E. coli* and *Enterococci* are commonly used to assess water quality (Holcomb and Stewart, 2020; USEPA, 2012). While FIB tests are convenient and relatively inexpensive, the test results do not necessarily indicate the recent occurrence of fecal contamination because some FIB can survive for long time and even grow in environments (Byappanahalli et al., 2012; Ishii and Sadowsky, 2008; Jang et al., 2017). Poor correlations were also reported between the levels of FIB and the occurrence of human pathogens (Harwood et al., 2005; Ishii et al., 2014b; Korajkic et al., 2018; Lemarchand and Lebaron, 2003; Oster et al., 2014; Wu et al., 2011), and as such, are also ineffective predictors of human health outcomes (Colford et al., 2007; Holcomb and Stewart, 2020). In addition, because FIB are largely ubiquitous in the digestive tracts of most warm-blooded animals, the levels of FIB alone cannot be used to elucidate sources of fecal pollution, making the regulation and remediation of pathogen-impaired waters a challenge.

Microbial source tracking (MST) is an approach to identify host-specific fecal microorganisms and pinpoint major sources of fecal contamination (and thus pathogens) in the environment (Harwood et al., 2014). Early applications of MST methods were primarily employed to discriminate between human fecal sources and other animal fecal sources, as human sewage was expected to attribute greater risks to the environment (Harwood et al., 2014). It is now understood that non-human fecal sources may contribute specific pathogens to the environment as well. For example, excrement from pigs may contain infectious Hepatitis E virus (Aggarwal and Naik, 2009). Wild and domestic avian species are known to excrete *Campylobacter* spp. (Kobayashi et al., 2022; Silva et al., 2011). Ruminants (i.e., cattle, sheep) are also known to be significant reservoirs of Shiga-toxin producing *E. coli* (Ishii et al., 2007; Paton and Paton, 1998). MST methods have expanded significantly in recent years in attempts to attribute pathogen loading to the correct fecal host. This methodology is being applied to new developments in Total Maximum Daily Load (Goodwin et al., 2017). However, the relationship between various MST markers and pathogens is still not well understood (Ahmed et al., 2013; Hughes et al., 2017; Korajkic et al., 2018), most likely because of the lack of comprehensive datasets of pathogens and MST marker concentrations.

High-throughput quantitative PCR is a powerful method to quantify multiple genes of interest for many samples (Ishii, 2019). When microfluidic technology is used to dispense and mix reagents and samples for high-throughput qPCR, it is called microfluidic qPCR (MFQPCR). We previously applied the MFQPCR technology (Fluidigm BioMark) to simultaneously quantify various pathogens and FIB in multiple samples (Ishii et al., 2014a; Ishii et al., 2013; Zhang and Ishii, 2018); however, these MFQPCR chips did not include assays for MST markers. Recently, Shahraki et al. (Shahraki et al., 2019) reported an MFQPCR OpenArray chip to simultaneously quantify bacterial pathogens, FIB, and MST markers. However, their chip includes a limited number of assays (15 bacterial pathogens, two FIB, and seven MST markers), and therefore, may not provide comprehensive data enough to analyze correlations between pathogen and MST marker concentrations.

Consequently, the objectives of this study were to (i) develop MFQPCR chips to simultaneously quantify pathogens (bacteria, viruses, and protozoa), FIB, and MST markers, (ii) apply the chips to quantify target organisms in wastewater, surface water, and fecal samples, and (iii) analyze correlations between pathogen and MST marker concentrations.

## 2. Materials and Methods

### 2.1. Quantitative PCR assays

A total of 80 TaqMan probe assays were selected from previous studies to be used in this research (**Table S1, S2, and S3**), including those for two FIB, 22 bacterial pathogens, 11 viral pathogens, five protozoan pathogens, 37 MST markers for various host species (general, human, dog, cow, pig, ruminant, avian, poultry, gull, goose, deer, beaver, muskrat), and three process controls. These assays demonstrated high sensitivity and specificity to the target organism, and have primer annealing temperatures of around 60°C. These assays were divided into two groups, DNA targets (Bacteria, Eukaryotes, and DNA viruses) and RNA targets (RNA viruses), and were run on separate MFQPCR chips.

All primers were synthesized and purified through a cartridge by Eurofins Genomics. All TaqMan probes were synthesized by Integrated DNA Technologies with 6-carboxyfluorescein (6-FAM) at their 5’ ends regardless of the fluorophore used in the original literature. When probes in the original literature were labeled with minor glove binder (MGB) at their 3’ ends, they were synthesized with MGB and a nonfluorescent quencher at their 3’ ends. When probes in the original literature were not labeled with MGB, they were synthesized with Iowa Black fluorescent quencher and an internal ZEN quencher that was inserted between the 9th and 10th bases from their 5’ ends. The addition of an internal ZEN quencher was previously shown useful to reduce the background fluorescent signals in TaqMan qPCR assays with non-MGB probes (Ishii et al., 2014a).

The gBlock DNA fragments containing target gene sequences were synthesized by Integrated DNA Technologies and used to generate the standard curves for qPCR (**Table S4**). The gBlocks DNA solutions (10^9^ copies/μl) for all 80 assays were pooled together and serially diluted to make 2 × 10^6^ – 2 × 10^0^ copies/μl of the standard DNA solutions.

### 2.2. Surface water samples

Surface water samples were collected from two areas of southwestern Lake Superior (46°43’N, 92°00’N) near the Duluth-Superior Harbor: Rice Point public water access and the Western Lake Superior Sanitary District (WLSSD) wastewater outfall point in St. Louis River, six times between June and October 2020 (**Fig. S1**). Rice Point is a popular boat launch used to enter both Lake Superior and the St. Louis River. Rice Point is adjacent (~0.4 km distance) to the Interstate Island State Wildlife Management Area, where thousands of common terns and ring-billed gulls nest each year. The Duluth Wastewater Outfall point is located approximately 1.1 km from Interstate Island.

To collect a sample at Rice Point, autoclave-sterilized L/S 18 precision pump tubing (Cole-Parmer) was weighted with an autoclave-sterilized hose barb and lowered into the water at the end of the dock (*ca*. 30 cm below the water surface). With a battery-powered peristaltic pump, approximately 40 L of water was pumped through a dead-end hollow-fiber REXEED 25S ultrafiltration membrane (Asahi Kasei) as described previously (Smith and Hill, 2009). After filtration, the ultrafiltration membrane was capped and transported on ice back to the laboratory, where the filter was backflushed within 48 h.

To collect samples from the Duluth wastewater outfall point, a canoe was launched from the Rice Point dock and paddled to the outfall location. Water sample (10 L) was collected 30 cm below the surface of the water as described above.

At both sites, water pH, conductivity, and temperature were measured by using the YSI Professional Plus Multiparameter Instrument (YSI, Yellow Springs, OH). Turbidity was measured by the Hach 2100P ISO portable turbidimeter (Hach, Loveland, CO). Upon returning to the lab (<12 h), total coliform and *E. coli* concentrations were measured using the Colilert test kits (IDEXX Laboratories).

### 2.3. Domestic wastewater samples

From November 2021 to May 2022, domestic wastewater influent and effluent samples were received from 11 wastewater treatment facilities throughout the United States. These facilities were geographically diverse, used various treatment strategies, and treated varied volumes of daily inflow. The locations and specific characteristics of participating facilities will remain anonymous, and samples are herein labeled as Facility “A” - “K”. Each facility collected a 10-L grab sample of finished effluent and a 1-L grab sample of raw influent. These samples were placed in a cooler on ice and shipped overnight to the laboratory at the University of Minnesota, St. Paul, MN, where they were processed within 48 h of collection. One effluent sample was unusable for filtration due to substantial leaking during shipment. To capture biomass from the effluent samples received, 10 L of effluent was pumped through a dead-end hollow-fiber REXEED 25S ultrafiltration membrane as described above. For influent samples, the filtration step was omitted. Approximately 700 mL of well-mixed influent was poured into a 1-L glass bottle and received the post-backflushing treatment as described below.

### 2.4. Water sample processing

Biomass was retrieved from the ultrafiltration membranes by backflushing with 500 ml of autoclave-sterilized backflushing solution (0.5% Tween 80, 0.01% sodium polyphosphate, and 0.001% Y-30 antifoam emulsion). Each backflushed product and unfiltered wastewater influent was mixed with 8% (w/v) polyethylene glycol (PEG) 8000, 1.15% (w/v) sodium chloride, and 1% sterile beef extract on a stir plate at 5°C for 1 h. After incubation at 5°C overnight, the treated backflushed product was transferred to 250 mL conical centrifuge bottles and then centrifuged at 5000 rpm and at 5°C for 45 min. Following centrifugation, the supernatant was decanted, and the remaining solids were suspended in 2 to 8 mL of 10X TE buffer (100 mM Tris, 10 mM EDTA [pH 8.0]). The final volume and mass of the suspension were measured prior to pipetting sample aliquots. The sample was thoroughly mixed, and six, 230 μL aliquots were pipetted into sterile microcentrifuge tubes for each sample. One set of triplicates (three of six samples) was immediately frozen at −80°C until DNA/RNA extraction. The remaining set received a propidium monoazide (PMA) dye treatment for live/dead cell distinction (see below).

To assess any background contamination from the ultrafiltration membranes or backflushing procedure, two blank filters were backflushed with 500 mL of sterile backflushing solution. Subsequent flocculation and centrifugation steps to concentrate biomass were performed as described above.

### 2.5. Fecal sample collection

Fresh gull and goose fecal samples were collected using sterilized plastic scoops from beaches in Duluth, MN, in August 2021. Samples were immediately placed into 15 mL tubes, kept on ice, and transported back to the laboratory. They were stored at – 80°C until DNA extraction. A total of ten gull and nine goose fecal samples were collected.

### 2.6. PMA Dye Treatment

To differentiate between viable and non-viable cells and viruses in qPCR, half of the aliquots mentioned above (i.e., the second set of triplicates) received a PMAxx dye treatment (Biotium, Fremont, CA). Aliquots of the biomass samples (230 μL) were mixed with 5.75 μL of 2 mM PMAxx dye (final PMAxx conc. of 50 μM) and incubated at room temperature in a dark box for 10 min, then exposed to light for 20 min in the PMA-Lite LED Photolysis Device (Biotium). After treatment, these samples were stored at – 80°C until DNA/RNA extractions.

### 2.7. DNA and RNA extractions

DNA and RNA were co-extracted from the triplicate biomass samples (PMA-treated and non-treated) originating from surface water, wastewater, and blanks, by using the AllPrep PowerViral DNA/RNA kit (Qiagen). Prior to bead-beating, 100 μL of phenol:chloroform:isoamyl alcohol (25:24:1, pH 6.6) and 6 μL of β-mercaptoethanol were added to extraction tubes for RNA isolation. Following this step, the extraction was performed following the manufacturer-recommended protocol. Resultant DNA/RNA solutions (100 μL) were stored at −80°C.

DNA was extracted from gull and goose fecal samples by using the QIAamp PowerFecal Pro DNA Kit (Qiagen) according to the manufacturer’s protocol. Resultant DNA solutions (100 μL) were stored at −80°C.

### 2.8. Reverse Transcription (RT) Reaction

To synthesize cDNA from viral RNA, reverse transcription was performed by using the PrimeScript RT Kit (Takara Bio, San Jose, CA) according to the manufacturer’s protocol. The RT reaction (10 μL) was done with 2.5 μM oligo dT primer, 20 μM random 6mers, and 2 μL of 10-fold diluted RNA samples. The RT product was stored at −20°C.

### 2.9. Conventional qPCR

Conventional qPCR was used to quantify select MST markers. The reaction mixture (10 μL) contained 1x SsoAdvanced Universal Probe Supermix (BioRad), 0.4 μM each of forward and reverse primers, 0.2 μM probe, and 1 μL DNA solution. Standard DNA (2 × 10^6^ – 2 × 10^0^ copies/μL) and no template control (NTC) were included in each run. The qPCR was done using StepOnePlus Real-Time PCR System (Applied Biosystems) under the following thermal conditions: initial denaturation at 98°C for 3 min, 40 cycles of 98°C for 10 s and 60°C for 30 s. ROX was used as a passive reference dye.

### 2.10. MFQPCR

Prior to MFQPCR, specific target amplification (STA) was done to increase the number of target molecules in the samples and standards. The STA reaction mixture (8 μL) contained 2× Prelude PreAmp master mix (Takarabio), 0.2 μM each primer, 1 μL of the gBlock solution (2 x 10^4^ copies/μL) for HF183-IAC assay, and 2 μL of the 10-fold diluted DNA samples or undiluted cDNA samples. The HF183-IAC gBlock solution was added as an internal amplification control (Green et al., 2014a). The STA reaction was done using a Veriti 96-Well Thermal Cycler (Applied Biosystems) with the following protocol: 95°C for 10 min, followed by 14 cycles with 95°C for 15 s and 60°C for 4 min. Upon completion, the STA product was diluted 5-folds by mixing with 32 μL TE buffer (10 mM Tris, 0.1 mM EDTA [pH 8.0]) and stored at −20°C.

To confirm the success of target amplification and assess any substantial amplification bias in the STA reaction, conventional qPCR was performed with pre- and post-STA standards as described above. STA reactions were deemed successful if the Ct values for the 50-fold diluted post-STA standards were approximately 4–8 times smaller than the Ct values for pre-STA standards.

MFQPCR was done using the BioMark HD Real-Time PCR system (Fluidigm) with the 96.96 DynamicArray IFC or 192.24 DynamicArray IFC for DNA or RNA targets, respectively. Aliquots (5 μL) of 10X assay mix (1X Assay Loading Reagent [Fluidigm], 2 μM each primer, 1 μM probe) and the sample mix (1X SsoAdvanced Universal Probe Supermix [BioRad], 1X Loading Reagent [Fluidigm], and 2.25 μl of the 5-fold diluted STA product) were loaded into the 96.96 or 192.24 chip and mixed using an IFC controller MX or RX (Fluidigm) for the 96.96 chip or the 192.24 chip, respectively, according to the manufacturer’s instructions. The final primer and probe concentrations after mixing were 200 nM and 100 nM, respectively. Standard DNA (2 × 10^6^ – 2 × 10^0^ copies/μL) and NTC were included in each run. To fill the 96.96 and 192.24 chips, some assays were run in duplicates. The qPCR was done under the following conditions: thermal mixing at 60°C for 30 s (only for the 96.96 chip), 98°C for 3 min, followed by 40 cycles of 98°C for 10 s and 60°C for 30 s. ROX was used as a passive reference dye.

### 2.11. Data Analysis

The MFQPCR results were analyzed using the Fluidigm Real-Time PCR Analysis software version 4.8.1. Threshold fluorescence intensity was manually set for each assay based on the logarithmic view of the amplification curve to obtain quantification cycle (Cq) values. Data were then exported to Microsoft Excel, where qPCR standard curves were generated based on the Cq value of the serially diluted standard DNA. Amplification efficiency (*E*) was calculated based on the slope of the standard curves with the following equation: *E* = −1 + 10^(1/slope)^ (Bustin et al., 2009). For environmental samples, the quantity of a target gene was calculated from the Cq value by using the standard curve.

Samples were considered quantifiable if the measured quantity was greater than the observable quantity in the lowest concentration standard (i.e., limit of quantification; LOQ) and the background signals in the NTC (if any). Samples that exhibited amplification but did not meet these criteria were considered Detected but Not Quantifiable (DNQ). Samples were considered Non-Detects (ND) for an assay if no amplification was observed within 40 cycles. Duplicate quantities (i.e., technical replicates) were averaged to obtain one value per sample.

We ran MFQPCR with three biological replicates (i.e., triplicate DNA samples per water sample). To obtain a single quantity per water sample, observed quantities were averaged across triplicate DNA samples. If two out of three replicates were quantifiable, the sample was considered quantifiable, and the mean gene quantity and standard error (SE) were calculated. If two out of three replicates were DNQ or if only one out of three replicates was quantifiable and another replicate was DNQ, the sample was considered DNQ. If samples did not meet these criteria, they were designated as ND.

Gene copy number per liter of water was calculated using the gene quantity value (copies/μL DNA), the mass of biomass pellet used for DNA/RNA extraction, the concentrated pellet mass, and the filtration volume for the water sample. For wastewater influent samples that omitted the filtration step, the filtration volume was substituted with the volume used for subsequent flocculation steps. Log reduction of microbes during wastewater treatment processes was calculated as follows: log reduction value (LRV) = log_10_ (influent gene concentration / effluent gene concentration).

Kendall’s rank correlations were analyzed between gene copy numbers and water quality parameters by using *corrplot* package in R (Wei and Simko, 2021). The correlation tests were performed with Bonferroni-adjusted p-values by using the *psych* package (Revelle, 2022). Only assays that were detected ≥50% of the samples were used for the correlation analysis. In addition, a log gene quantity was randomly imputed for each of the DNQ and ND samples by using the NADA package (Lee, 2020) and the Monte Carlo simulation with a sample size of 1×10^6^. To estimate the values for DNQ samples, the mean and standard deviation were estimated for each assay by using the *cenfit* function and the assay-specific limit of quantification (**Table S5**). Similarly, the values for ND samples were randomly imputed using the mean and standard deviation estimated based on the limit of detection for each assay, which is the log gene quantity at Cq = 40 calculated using each assay’s standard curve.

## 3. Results

### 3.1. Impacts of specific target amplification on qPCR performance

Specific target amplification (STA) was done to increase the number of target molecules in the samples and standards. The impact of STA was examined by conventional qPCR with six randomly selected assays (H8, Gull-4, Entero1, EV, HNoV-GI, Ehf) (**Fig. 1**). These assays demonstrated amplification efficiencies ranging from 78% – 104% and the LOQ as low as 2 – 20 gene copies/μL. These values were similar between pre-STA and post-STA standards. The Ct values for post-STA standards were four to nine times smaller than those for pre-STA standards. Considering the 1/50 dilution of the STA product prior to qPCR, the preamplification of samples increased the target DNA molecules by approximately 3 × 10^3^ – 2 × 10^5^ times.

**Figure 1.**
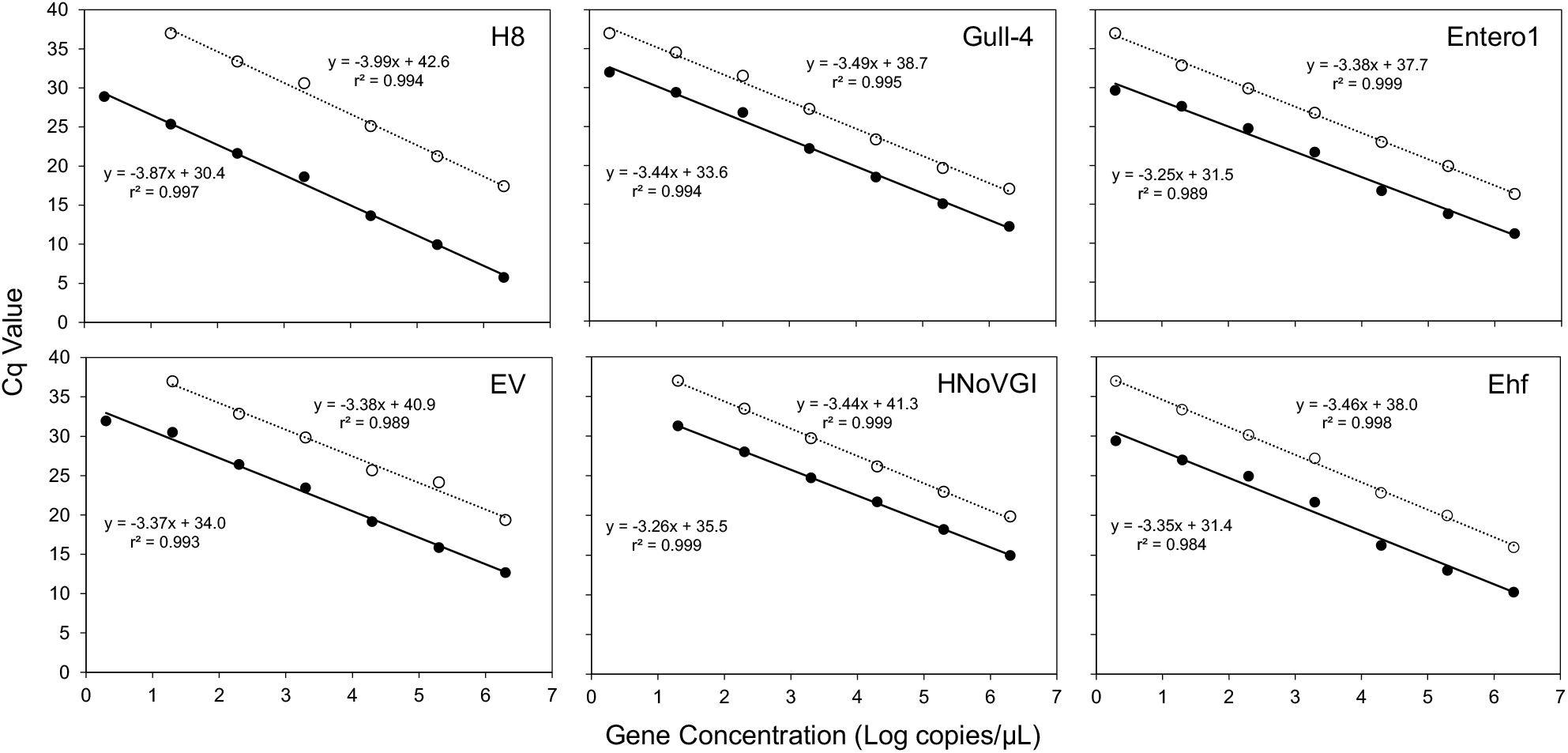
PCR standard curves for six, randomly selected assays for STA method validation. The standard curves were created by linear regression analysis of reaction Cq value versus the known concentration of standard DNA (log copies/μL). The two standard curves are shown per assay: one with pre-STA (○) and another with post-STA standards (●). Regression equations for the standard curve and r^2^ values are shown.

### 3.2. Analytical sensitivity and efficiency of MFQPCR assays

Out of the 80 assays included in the MFQPCR platform, 76 assays showed consistent amplification over multiple chip runs. Four assays (BacCow-UCD, Gull2Taqman, HNoV-GII, LA35) showed an absence of or inconsistent standard DNA amplification, and therefore, were removed from the downstream analyses. For the 76 assays, r^2^ values of the standard curves were >0.97 and the LOQ ranged from 2 to 2000 copies/μL (**Table S2**). The LOQ of the assay used to detect human adenovirus (ADV) was high because of the background signals in NTC. Amplification efficiencies of the 76 assays in the MFQPCR platform ranged from 71% to 125%. According to the MIQE guidelines, the generally accepted range for qPCR amplification efficiency is 80% – 120% (Bustin et al., 2009). Three assays (*gyrB*, H8, and MuBa01) fell below the acceptable range, with efficiencies of 71% – 79%, while one assay (HF183-IAC) exhibited an efficiency of 125%, slightly above the acceptable threshold.

### 3.3. Specificity of the MFQPCR assays

The high specificity of the MST assays used in this study was previously examined using conventional qPCR. This study confirmed their specificity, especially of human MST markers, on the MFQPCR platform. Human MST markers were detected in >81% of the influent wastewater samples but not from goose and gull fecal samples (**Fig. 2**). Among the human MST markers, crAssphage markers (CPQ_056 and CPQ_064 assays) were detected most frequently (100%) in the influent wastewater samples. Some non-human markers, especially those for dogs (DogBact and BacCan-UCD assays), cows (Cow-ND5 and BacBov1 markers), and muskrats (MuBa01 assay) were also detected in wastewater samples.

**Figure 2.**
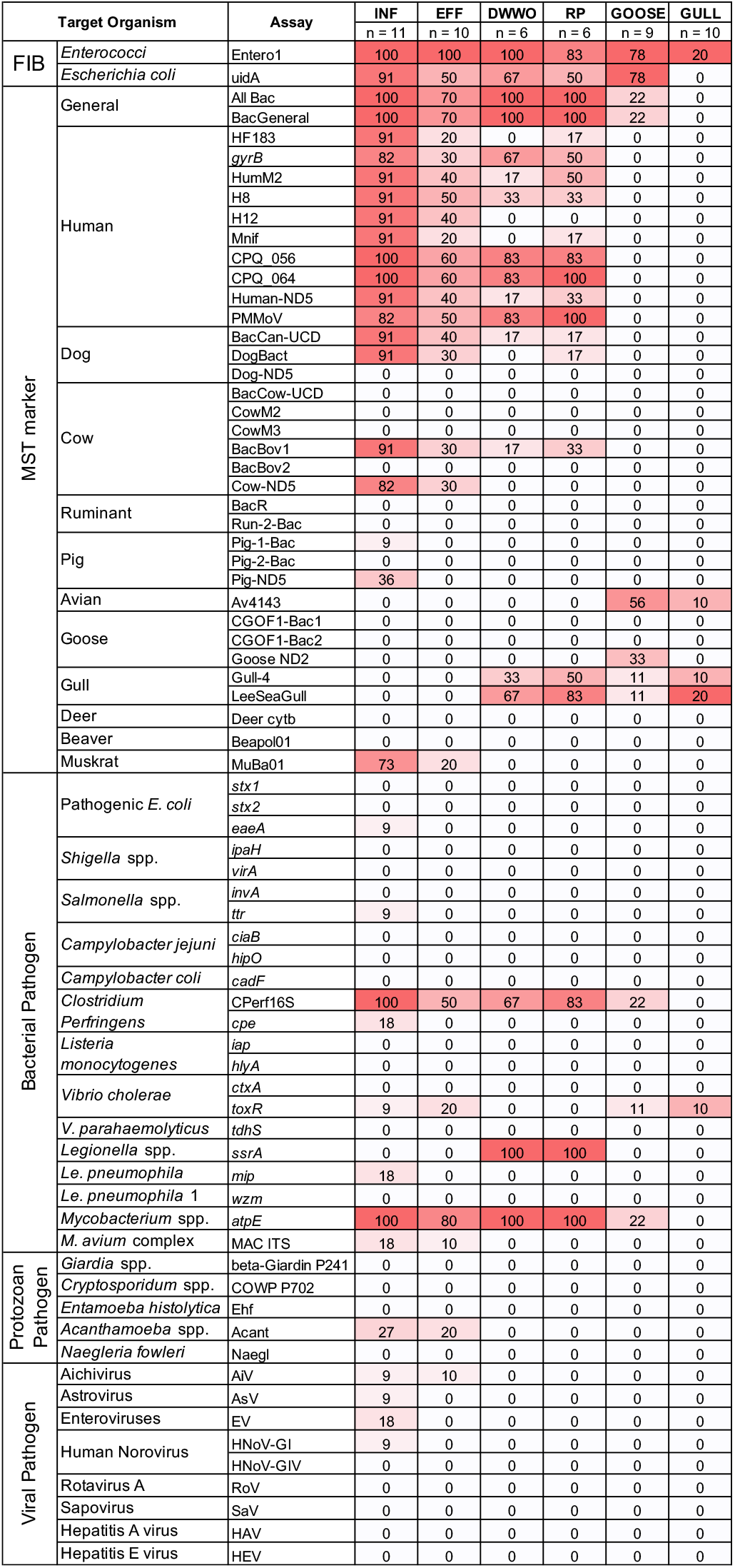
Proportion of quantifiable samples per sample and assay type. Samples were considered quantifiable if the quantity detected was greater than the Limit of Quantification (LOQ). Legend: INF, wastewater influent; EFF: wastewater effluent; DWWO, Duluth wastewater outfall; RP, Rice Point; GOOSE, goose fecal samples; GULL; gull fecal samples.

The MST markers for avian (Av4143 assay) and goose (CGOF1-ND2) were detected only in the avian (goose/gull) and goose feces, respectively, while gull markers (Gull-4 and LeeSeaGull assays) were detected in goose and gull feces. Their frequency of detection in the target organisms was relatively low (10–55%). Other goose markers (CGOF1-Bac1 and CGOF1-Bac2 assays) were not detected in the goose and gull feces collected in this study.

### 3.4. Internal amplification and blank controls

The internal amplification control (HF183-IAC assays) was detected in all samples at similar Cq values (13.05 ± 0.33 SD). The lack of false negatives and the consistent quantification of the IAC in all samples suggest that the PCR inhibition by potential DNA impurities was negligible. In addition, blank filters did not show quantifiable detection of genes, suggesting minimal background contamination occurred during sample processing.

### 3.5. Quantification of pathogens, FIB, and MST markers in wastewater and surface water samples

Various pathogens, FIB, and MST markers were quantitatively detected by MFQPCR in wastewater and surface water samples (**Fig. 2**). Fecal indicators *Enterococcus* spp. (Entero1 assay) and *E. coli (uidA* assay) as well as general *Bacteroides* markers (All Bac and BacGen assays) were frequently (50–100%) detected, while most bacterial, viral, and protozoan pathogens were detected relatively infrequently (<30%) across all samples. Exceptions to this include *Clostridium perfringens* (CPerf16S assay) which were detected in 67–100% of wastewater and surface water samples, and *Mycobacterium* spp. (*atpE* assay), which were detected in 80–100% of wastewater and surface water samples. Additionally, *Legionella* spp. (*ssrA* assay) was detected in 83–100% of Rice Point and Duluth Wastewater Outfall samples.

The MFQPCR also provided quantitative information. Concentrations of FIB and MST markers were generally higher than those of pathogens (**Fig. 3**). In addition, the gene concentrations of PMA-treated samples were lower than those of non-treated samples across all assays (*p* <0.05 by ANOVA). This is likely due to the amplification from dead/damaged cells was suppressed by the PMA treatment.

**Figure 3.**
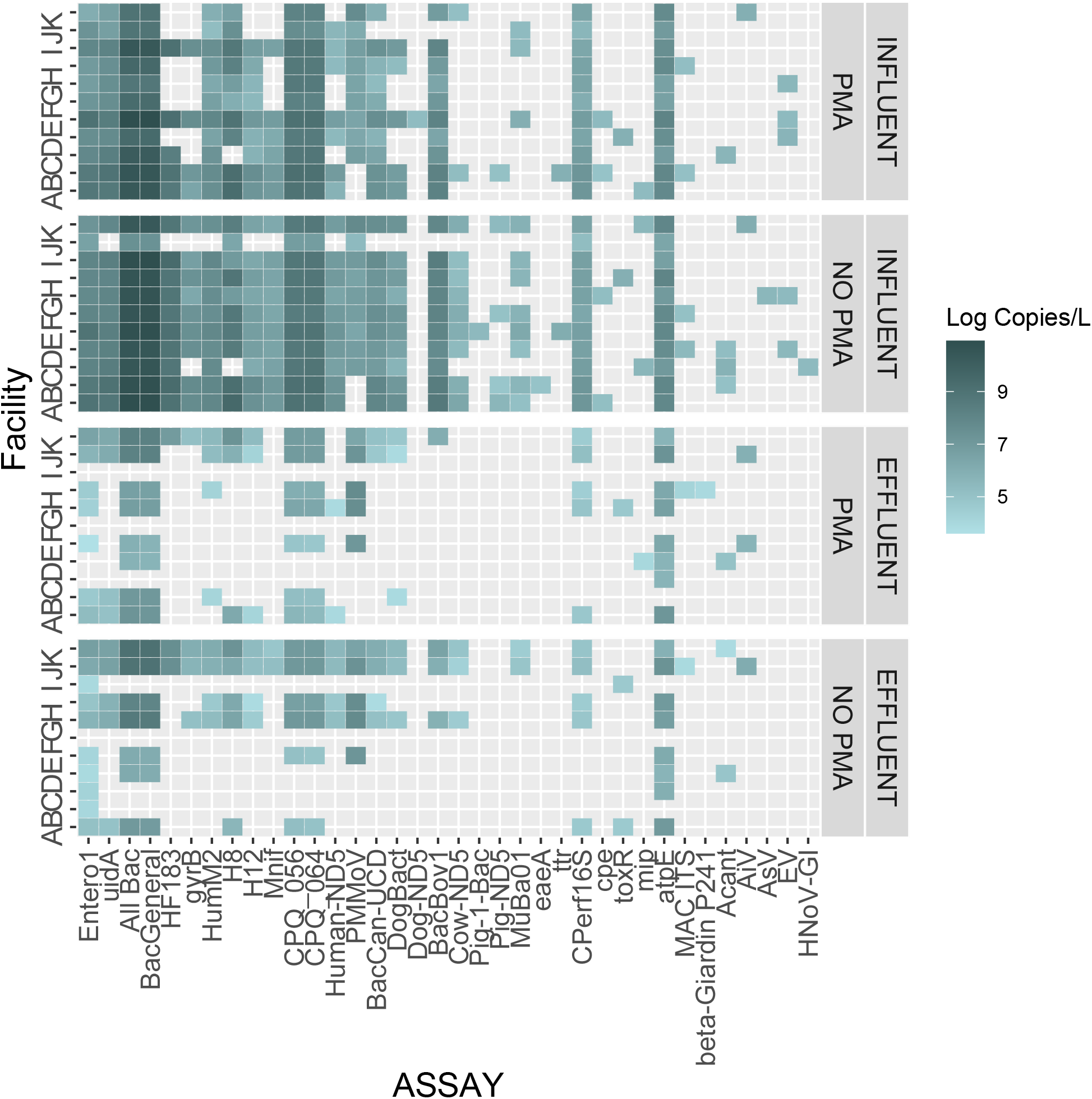
Heatmap of log gene copies per liter of wastewater influent and effluent from Facilities A-K. Gene quantification was done using MFQPCR with and without PMA treatment.

The concentrations of various bacteria and crAssphage in wastewater decreased, on average, by about 1-2 logs after treatment, although the log reduction value (LRV) varied by assays (**Table 1**; *p* <0.001 by ANOVA). The LRV for PMMoV was negative for both PMA-treated and non-treated samples (−0.53 and −0.72, respectively), indicating that their concentration was greater in effluent than in influent.

The gene concentration patterns were similar between the Rice Point and Duluth Wastewater Outfall samples (**Fig. 4**). The concentrations of some human MST markers (CPQ_056, CPQ_064, and Human-ND5 assays) were higher in the outfall samples than in the Rice Point samples. In contrast, the concentrations of gull MST markers (LeeSeaGull and Gull-4 assays) were higher in the Rice Point samples than in the outfall sample.

**Figure 4.**
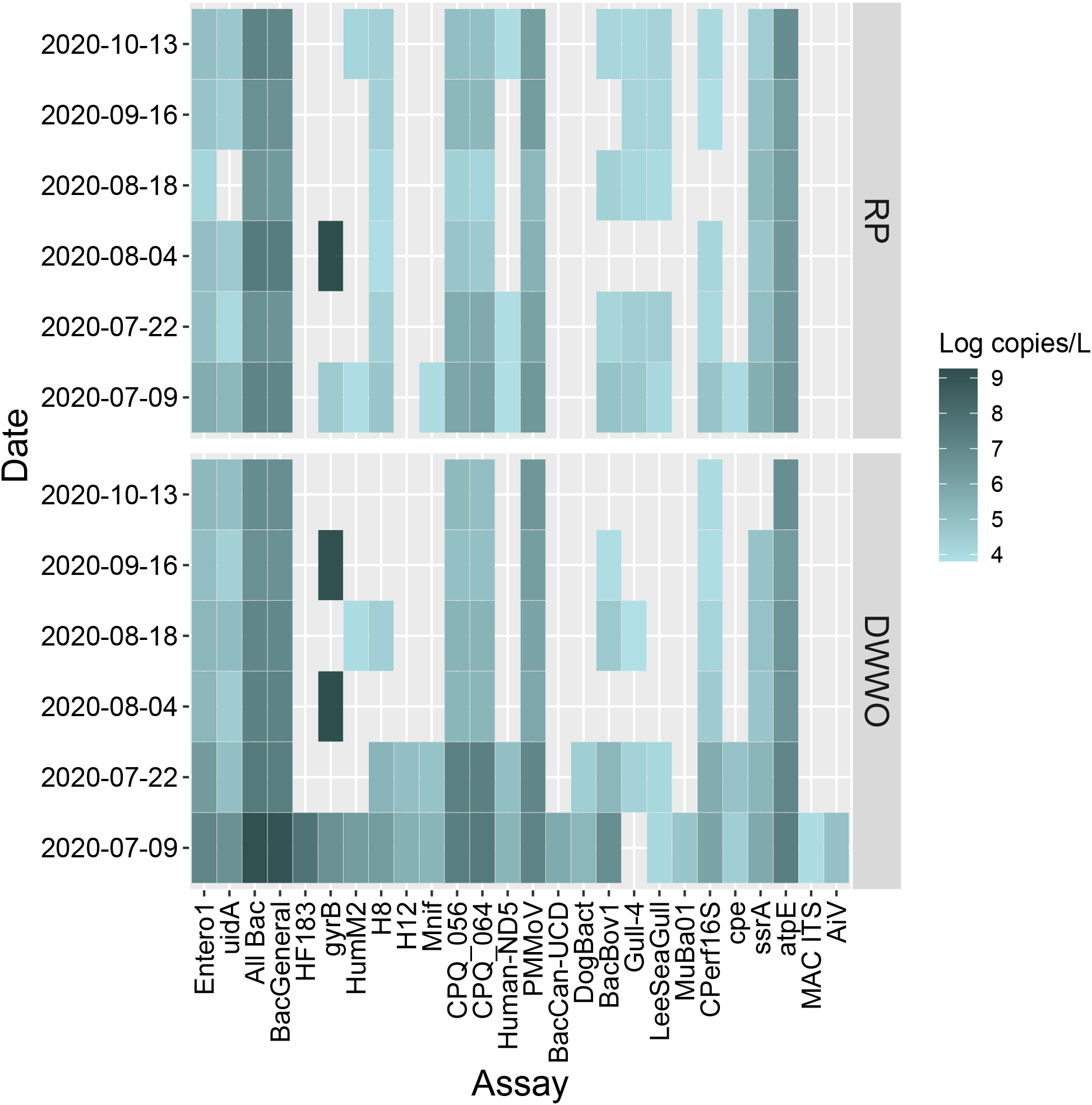
Concentrations of viable FIB, MST markers, and pathogens in surface water samples. Biomass from the water samples was treated with PMA to make the DNA from dead cells unavailable for PCR. The results of PMA-untreated samples are shown in Figure S2. Legend: RP, Rice Point; DWWO, Duluth wastewater outfall.

### 3.6. Correlations between pathogen, FIB, and MST marker concentrations

Correlation analysis was done with the PMA-treated surface water samples. Positive correlations (tau > 0.4) were seen between most of the human MST markers (**Fig. 5**), some of which were significantly correlated (**Table S6**). A positive correlation (tau = 0.48) was also seen between gull markers (Gull_4 vs. LeeSeaGull), although this was not statistically significant. Concentrations for FIB (uidA and Entero1 assays), general *Bacteroides* (All Bac and BacGeneral assays), *Clostridium perfringens* (CPerf16S assay), and *Mycobacterium* spp. (*atpE* assay) were also positively correlated with many of the human MST markers; whereas the concentration for *Legionella* spp. (*ssrA* assay) were significantly and positively correlated with cow MST marker (BacBov1; tau = 0.64, *p* = 0.026). Correlations were also seen between human MST markers and water quality parameters (e.g., water temperature, turbidity, and conductivity).

**Figure 5.**
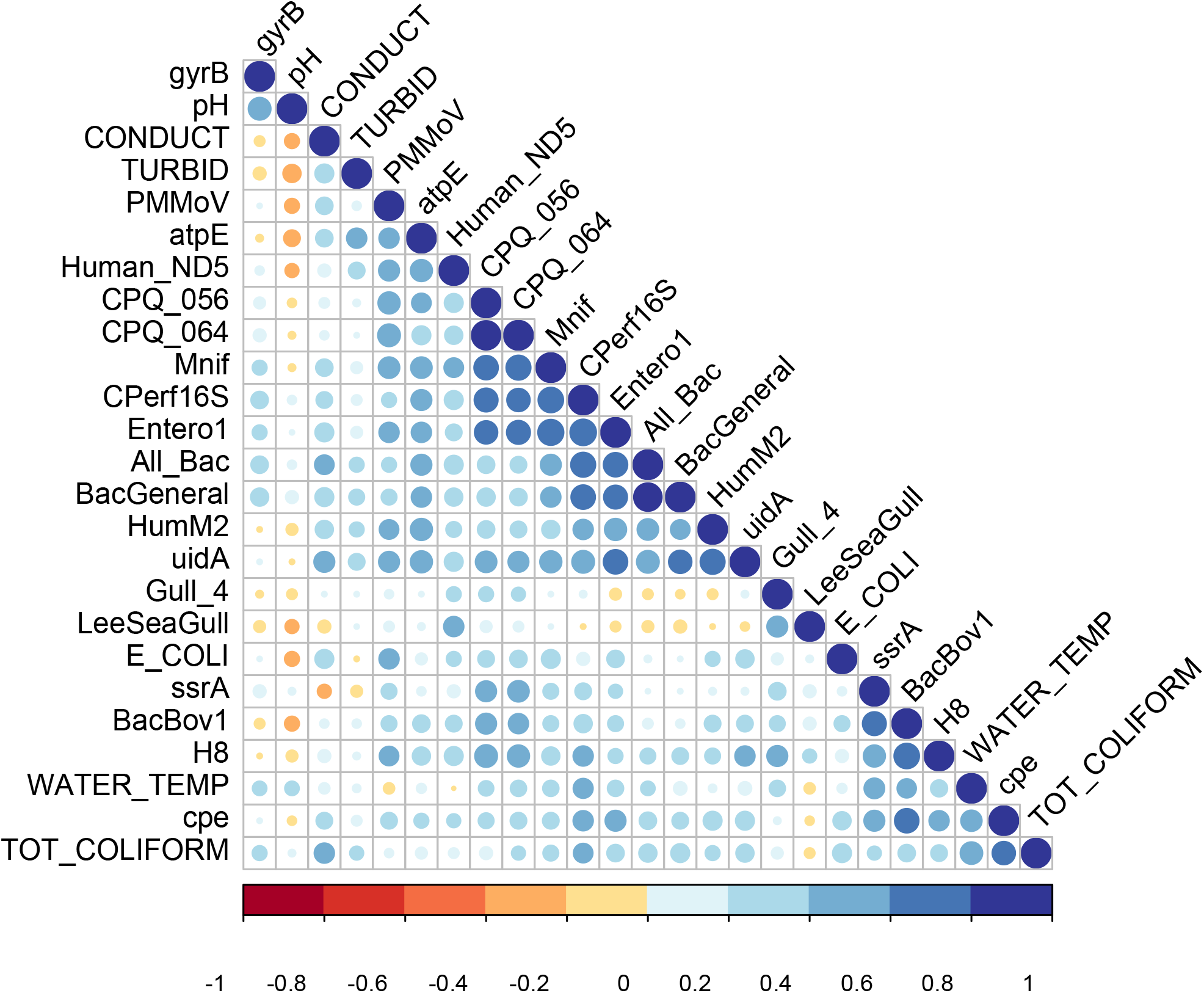
Correlation plot showing the associations between MST markers, FIB, and pathogen concentrations, and water quality parameters. This plot was generated using a matrix of Kendall’s Tau correlation coefficients. Statistical significance values of these correlations are shown in Table S6.

## 4. Discussion

High-throughput qPCR, including MFQPCR, has been shown useful to quantify genes of interest in environmental samples. We previously reported MFQPCR chips to simultaneously quantify various bacterial (Ishii et al., 2013; Zhang and Ishii, 2018) and viral pathogens (Ishii et al., 2014a); however, MST markers were not included on these chips. In addition, the TaqMan probes reported in Ishii et al. (Ishii et al., 2013) are no longer available; therefore, we needed to re-establish a new chip format. We updated our MFQPCR chip format by adding MST markers (37 assays), increasing the number of pathogens targeted (22, 11, and 5 assays for bacterial, viral, and protozoan pathogens, respectively), and adding internal amplification and process controls (three assays) as well as FIB (two assays). Recently, Shahraki et al. (Shahraki et al., 2019) reported a similar MFQPCR OpenArray chip to simultaneously quantify 15 bacterial pathogens, two FIB, and seven MST markers (a total of 24 target genes) for recreational water monitoring; however, the system developed in this study can provide more comprehensive data because it can quantify (i) both bacterial, viral, and protozoan pathogens, (ii) larger number of MST assays, (iii) internal amplification and process controls, and (iv) with more samples per run (up to 88 and 184 samples when using 96.96 and 192.24 chips, respectively, excluding standards and NTC).

Instead of designing new assays, we used previously validated TaqMan qPCR assays that are specific and sensitive to detect target organisms. The analytical sensitivity of these assays was also verified on the MFQPCR platform in this study, with LOQ of 2–20 copies/μL for most assays. The high analytical sensitivity on the MFQPCR platform was made possible by the STA reaction. The STA reaction is a multiplex PCR with all primers used for MFQPCR and with a small number of PCR cycles (10–14) (Ishii et al., 2013). Theoretically, at 100% amplification efficiency, a 14-cycle STA reaction should increase template DNA yield by 2^14^ times. The STA reaction is necessary to provide enough amount of template DNA to the MFQPCR reaction chambers (i.e., 6.7 nL) and allow for the detection of low-copy-number genes (Ishii et al., 2013). Similar to this study, STA reaction was previously used to increase template DNA molecules without major amplification biases (Ishii et al., 2014a; Ishii et al., 2013; Mathai et al., 2019; Sandberg et al., 2018; Zhang and Ishii, 2018). We did not examine the effects of STA for all assays, but based on the previous literature (Korenková et al., 2015), it is unlikely that our STA conditions caused significant biases in the amount of target DNA molecules.

The specificity of the MST assays used in this study was previously examined by using conventional qPCR (see references in **Table S1**). Therefore, we did not re-examine the specificity of the MST assays with a suite of host animals. Nonetheless, our MFQPCR results verified the specificity of the human MST assays. The human MST markers used in this study were frequently and abundantly detected in wastewater samples, but they were not detected from avian feces. Similarly, avian (goose and gull) markers occurred only in avian feces, although they were detected relatively infrequently and at low levels. The infrequent and low levels of these MST markers in the fecal samples may be related to microbial decay. An effort was made to select the freshest possible fecal samples; however, our samples may have included some aged feces, which could impact the analysis of MST marker presence for goose and gull samples.

Dog MST markers were also detected in wastewater samples similar to previous reports (Green et al., 2014b; Kildare et al., 2007; Schriewer et al., 2013). Although some dog MST markers (e.g., BacCan-UCD) are reported to cross-react with human feces (Kildare et al., 2007), it is also possible that wastewater actually contained both human and dog secreta (Green et al., 2014b). Similarly, some cow MST markers are also known to cross-react with human wastewater (Ahmed et al., 2013). Because most MST markers cross-react with non-target host DNA to a limited extent, it is important to analyze multiple assays for the same MST host (which was done on our MFQPCR chip format) to increase the accuracy of the source identification (Ahmed et al., 2019; Ahmed et al., 2013).

In addition to a suite of MST markers, our MFQPCR chips can also quantify various bacterial, viral, and protozoan pathogens. The detection frequencies and concentrations of these pathogens were much lower than those of MST markers similar to previous studies (Ahmed et al., 2013; Zhang et al., 2016). While *Legionella* spp. (*ssrA* assay), *Mycobacterium* spp. (*atpE* assay), and *Clostridium perfringens* (CPerf16S assay) were relatively frequently detected in wastewater and surface water samples, these bacteria include both pathogenic and non-pathogenic strains. Because the three assays mentioned above do not target virulence factor genes, the quantities of *Legionella* spp., *Mycobacterium* spp., and *Clostridium perfringens* measured by these assays do not necessarily indicate the level of human pathogens. Multiple species of *Legionella*, *Mycobacterium*, and *Clostridium* are known to be naturally occurring in aquatic environments (Atlas, 1999; Kiu and Hall, 2018; Mueller-Spitz et al., 2010; Primm et al., 2004). Assays that target virulence factor genes (e.g., cpe assay for *Clostridium perfringens* enterotoxin) or more pathogen-related subgroups (e.g., mip assay for *L. pneumophila*, wzm assay for *L. pneumophila* serogroup 1, and MAC ITS assay for *Mycobacterium avium* complex) could provide more pathogen-related information. The infrequent detection of these assays in this study suggested low pathogenic potential.

Quantitative information obtained by MFQPCR can be used to evaluate the log reduction of microbes during wastewater treatment processes. While the concentrations of most bacteria and crAssphage decreased by about 1-2 logs, that of PMMoV did not. Similarly, greater removal of crAssphage than PMMoV was previously reported (Tandukar et al., 2020), although the concentrations of both viruses decreased (3.3 ± 1.0 and 2.0 ± 0.4 for crAssphage and PMMoV, respectively). The different LRV between Tandukar et al. (Tandukar et al., 2020) and this study may be related to different treatment systems and/or water temperatures (Canh et al., 2019). It is also possible that different water sample processing between WWTP effluent and influent samples (i.e., with and without ultrafiltration, respectively) could have influenced the recovery of viral particles results in this study.

Pathogen concentrations measured by MFQPCR can be also useful for quantitative microbial risk assessment (QMRA). One of the criticisms associated with the qPCR-based QMRA is the overestimation of risk by detecting signals from dead cells and viral particles (Ishii et al., 2014b). PMA treatment has the potential to overcome this issue by reducing the qPCR signals from dead cells/particles (Nocker et al., 2007). Lower gene concentrations in the PMA-treated samples than in non-treated samples observed in this study suggest that PMA treatment successfully reduced the signals from dead cells/particles.

The MFQPCR chips developed in this study was also applied to identify potential sources of fecal contamination at two locations. While both human and gull MST markers were detected from the two sites, the human MST markers were more abundant in the Duluth wastewater outfall samples than in the Rice Point samples, suggesting that the outfall samples were more impacted by human sources than the Rice Point samples. In contrast, the concentrations of gull markers were higher in the Rice Point samples than the outfall sample. This was likely due to Rice Point’s proximity to the gull population on Interstate Island. Similar results were obtained by previous MST studies done at or near these sites (Eichmiller et al., 2013; Hansen et al., 2011).

Positive correlations between FIB, general *Bacteroides*, and many human MST markers suggested that the main source of fecal contamination was humans in the Duluth wastewater outfall and Rice Point samples. Similar results from multiple markers can increase the confidence of the source identification as suggested previously (Ahmed et al., 2019). Genes targeting *Clostridium perfringens* and *Mycobacterium* spp. were also positively correlated with human MST markers, suggesting that these potential human pathogens likely originated from humans in these water samples. In contrast, a positive correlation was seen between *Legionella* spp. *ssrA* and a cow MST marker BacBov1, suggesting that *Legionella* spp. may be originated from cows. This is also supported by the isolation of *Legionella* spp. from cattle (Boldur et al., 1987; Fabbi et al., 1998). However, since only one out of six cow MST markers showed this relationship, care should be taken when interpreting the result. For example, we cannot exclude the possibility of the BacBov1 assay detecting signals from other hosts (e.g., humans) similar to other cow MST markers (Ahmed et al., 2013). More samples need to be analyzed to clarify the relationship between *Legionella* and cow MST markers.

In this study, we examined simple correlations; however, multivariate regression or similar modeling approach can be used to predict the occurrence of pathogens in a given water sample based on the concentrations of MST markers, FIB, and/or additional physicochemical parameters (Hill, 2022). Such a modeling approach could expand the toolkit available for water quality monitoring and QMRA, and therefore, should be tested in the near future.

## 5. Conclusions

- The MFQPCR chips developed in this study can provide quantitative information on various pathogens, FIB, and MST markers in environmental water samples.
- Multiple assays are included for the same pathogen or host to increase the accuracy of the pathogen or source identification.
- The MFQPCR results are useful to analyze pathogen removal rates, assess human health risks, identify potential sources of fecal contamination, and examine correlations between pathogen and MST marker concentrations.
- The comprehensive dataset of pathogens, FIB, and MST markers can provide opportunities to generate predictive models to estimate the presence of pathogens in a given water sample based on the concentrations of MST markers, FIB, and/or additional physicochemical parameters.

## Supporting information

Supplementary Matrials

## CRediT authorship contribution statement

**Ellie Hill:** Conceptualization, Methodology, Formal analysis, Investigation, Writing – original draft, Visualization. **Chan Lan Chun:** Resources, Writing – review & editing, coordination, Supervision. **Kerry Hamilton:** Conceptualization, Resources, Writing – review & editing, Funding acquisition. **Satoshi Ishii:** Conceptualization, Methodology, Formal analysis, Resources, Writing – review & editing, Visualization, Supervision, Funding acquisition.

## Declaration of Competing Interest

The authors declare no conflict of interest.

## Acknowledgements

We thank John Griffith and Josh Steele (SCCWRP) and John Meschke (University of Washington) for assistance in connecting with local WWTP facilities. We also thank those participating anonymous wastewater treatment facilities for providing water samples to this research, and Tamara Walsky, Caitlin Graeber, Maxwell Brubaker, Collin Krochalk, and Braeden Cox for their work in sample collection and processing.

This work was supported by the National Science Foundation award CBET 1916025 and the MnDRIVE Initiative of the University of Minnesota. E.H. was supported by the Moos Graduate Research Fellowship in Aquatic Biology.

## Notes

### Competing Interest Statement

The authors have declared no competing interest.

