## Supplementary Matrials for "A High-Throughput Microfluidic Quantitative PCR Platform for the Simultaneous Quantification of Pathogens, Fecal Indicator Bacteria, and Microbial Source Tracking Markers"

**Figure S1.** Locations of Rice Point boat launch and WLSSD effluent discharge point (Duluth Wastewater Outfall) in Duluth, MN.
**Figure S2.** Concentrations of FIB, MST markers, and pathogens in surface water samples.
**Table S1.** Primer and probe sequences used in this study to quantify microbial source tracking markers (DNA targets).
**Table S2.** Primer and probe sequences used in this study to quantify fecal indicator bacteria, pathogenic bacteria, protozoans, DNA virus, and internal amplification controls (DNA targets)
**Table S3.** Primer and probe sequences used in this study to quantify RNA targets.
**Table S4.** gBlock sequences used as DNA standards.
**Table S5.** Amplification efficiency (E), limit of quantification (LOQ), and $r^2$ value of the MFQPCR assays.
**Table S6.** Statistical significance (p value) of Kendall's rank correlations between gene copy numbers and water quality parameters.

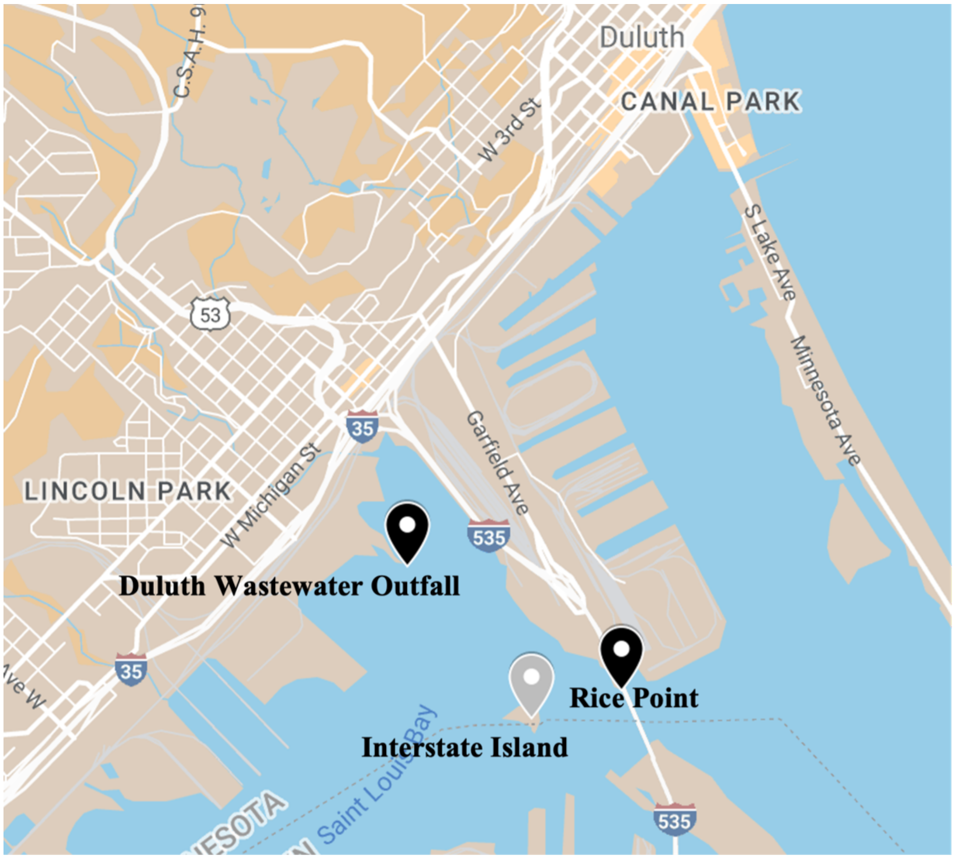


**Figure S1.** Locations of Rice Point boat launch and WLSSD effluent discharge point (Duluth Wastewater Outfall) in Duluth, MN. Interstate Island State Wildlife Management Area is labeled for reference. Figure generated using Google Map.

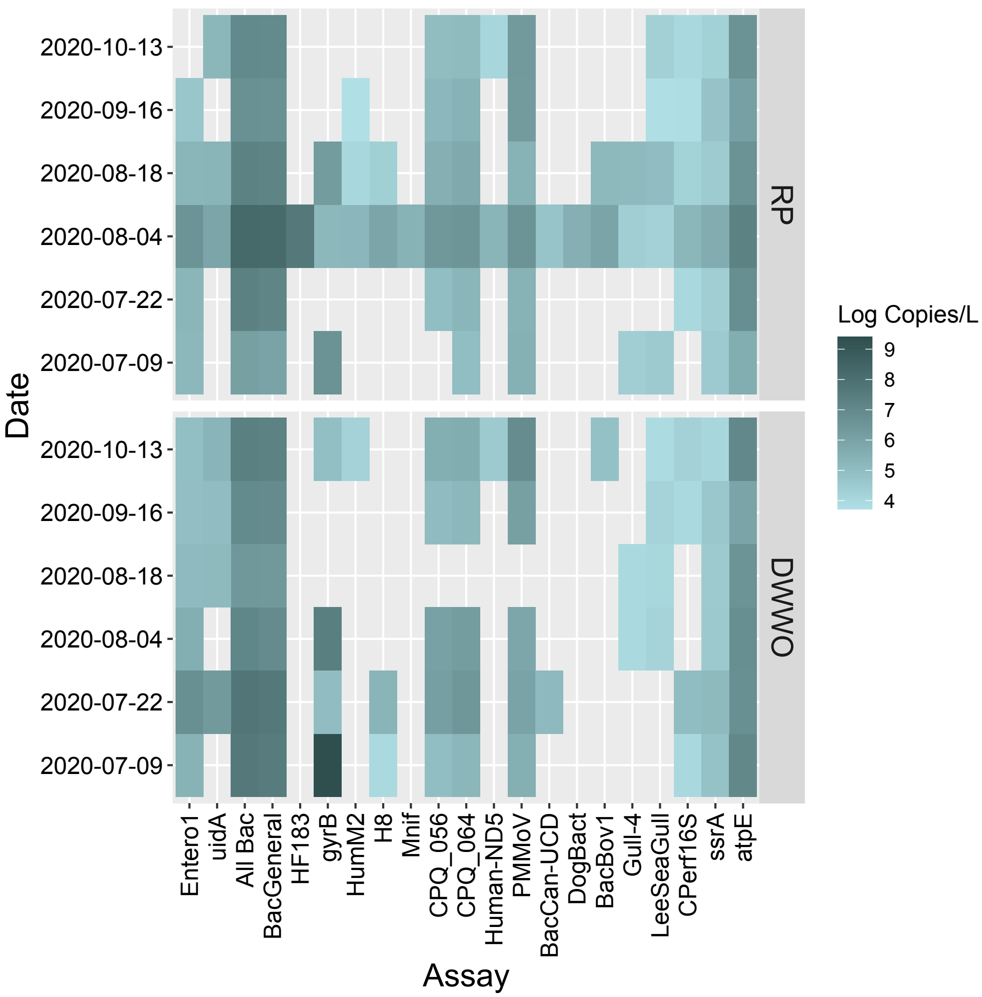


**Figure S2.** Concentrations of FIB, MST markers, and pathogens in surface water samples. Biomass from the water samples were not treated with PMA; therefore, the concentrations shown here represent both viable and dead cells. Results of PMA-treated samples are shown in Figure 4. Legend: RP, rice point; DWWO, Duluth wastewater outfall.

**Table S1.** Primer and probe sequences used in this study to quantify microbial source tracking markers (DNA targets).

| Target Organism | Assay Name | Primer/Probe Name | Primer sequence (5' --> 3') | Reference |
|---|---|---|---|---|
| General Bacteroides | | | | |
| *Bacteroides* spp. | AllBac | AllBac296f | GAGAGGAAGGTCCCCCAC | (1) |
| | | AllBac412r | CGCTACTTGGCTGGTTCAG | |
| | | AllBac375Bhqr | **FAM-**CCATTGACC/**ZEN/**AATATTCCTCACTGCTGCCT**-IBFQ** | |
| | BacGeneral | BacGen-F | CTGAGAGGAAGGTCCCCCAC | (2) |
| | | BacGen-R | CACGCTACTTGGCTGGTTCAG | |
| | | BacGen-TP | **FAM-**AGCAGTGAGGAATATT**-MGB-NFQ** | |
| Human | | | | |
| Human *Bacteroides* | HF183 | HF183 | ATCATGAGTTCACATGTCCG | (3) |
| | | BacR287 | CTTCCTCTCAGAACCCCTATCC | |
| | | BacP234MGB | **FAM-**CTAATGGAACGCATCCC**-MGB-NFQ** | |
| | *gyrB* | Bf904F | GGCGGTCTTCCGGGTAAA | (4) |
| | | Bf1272R | TGGCATATAGCGGAAGAAAAAAG | |
| | | Bf923MGB | **FAM-**TGGCCGACTGCTC**-MGB-NFQ** | |
| | HumM2 | Hum2F | CGTCAGGTTTGTTTCGGTATTG | (5) |
| | | Hum2R | TCATCACGTAACTTATTTATATGCATTAGC | |
| | | Hum2P | **FAM-**TATCGAAAA/**ZEN/**TCTCACGGATTAACTCTTGTGTACGC**-IBFQ** | |
| Human-specific *E. coli* | H8 | H8_Forward | ACAGTCAGCGAGATTCTTC | (6, 7) |
| | | H8_Reverse | GAACGTCAGCACCACCAA | |
| | | H8_Probe | **FAM-**ACTGGCATC/**ZEN/**GGCATGGAACAC**-IBFQ** | |
| | H12 | H12_Forward | GTAAAAGGACTGCCGGGAAA | (6, 7) |
| | | H12_Reverse | TCAGATCGTCCTTTACCAG | |
| | | H12_Probe | **FAM-**AGAGTAAAC/**ZEN/**GCTTCGCCCTTAGCC**-IBFQ** | |

**Table S1.** Continued.

| Target Organism | Assay Name | Primer/Probe Name | Primer sequence (5' --> 3') | Reference |
|---|---|---|---|---|
| *Methanobrevibacter smithii nifH* | Mnif | Mnif_202F | GAAAGCGGAGGTCCTGAA | (8) |
| | | Mnif_353R | ACTGAAAAACCTCCGCAAAC | |
| | | Mnif Probe | **FAM-**CCGGACGTG/**ZEN**/GTGTAACAGTAGCTA**-IBFQ** | |
| crAssphage | CPQ_056 | 056F1 | CAGAAGTACAAACTCCTAAAAAACGTAGAG | (9) |
| | | 056R1 | GATGACCAATAAACAAGCCATTAGC | |
| | | 056P1 | **FAM-**AATAACGATTTACGTGATGTAAC**-MGB-NFQ** | |
| | CPQ_064 | 064F1 | TGTATAGATGCTGCTGCAACTGTACTC | (9) |
| | | 064R1 | CGTTGTTTTCATCTTTATCTTGTCCAT | |
| | | 064P1 | **FAM-**CTGAAATTGTTCATAAGCAA**-MGB-NFQ** | |
| Human mtDNA | Human-ND5 | Human-ND5_forward | CAGCAGCCATTCAAGCAATGC | (10) |
| | | Human-ND5_reverse | GGTGGAGACCTAATTGGGCTGATTAG | |
| | | Human-ND5_probe | **FAM-**TATCGGCGA/**ZEN**/TATCGGTTTCATCCTCG**-IBFQ** | |
| Dog | | | | |
| Dog *Bacteroides* | BacCan-UCD | BacCan-545f1 | GGAGCGCAGACGGGTTTT | (11) |
| | | BacUni-690r1 | CAATCGGAGTTCTTCGTGATATCTA | |
| | | BacUni-690r2 | AATCGGAGTTCCTCGTGATATCTA | |
| | | BacUni-656p | **FAM-**TGGTGTAGCGGTGAAA**-MGB-NFQ** | |
| | DogBact | DF475F | CGCTTGTATGTACCGGTACG | (12, 13) |
| | | Bac708R | CAATCGGAGTTCTTCGTGAT | |
| | | DogBactP | **FAM-**ATTCGTGGT/**ZEN**/GTAGCGGTGAAATGCTTAG**-IBFQ** | |
| Dog mtDNA | Dog-ND5 | Dog-ND5_forward | GGCATGCCTTTCCTTACAGGATTC | (14) |
| | | Dog-ND5_reverse | GGGATGTGGCAACGAGTGTAATTATG | |
| | | Dog-ND5_probe | **FAM-**TCATCGAGT/**ZEN**/CCGCTAACACGTCGAAT**-IBFQ** | |

**Table S1.** Continued.

| Target Organism | Assay Name | Primer/Probe Name | Primer sequence (5' --> 3') | Reference |
|---|---|---|---|---|
| Cow | | | | |
| Cow *Bacteroides* | BacCow-UCD | BacCow-CF128f | CCAACYTTCCCGWTACTC | (11, 15) |
| | | BacCow-305r | GGACCGTGTCTCAGTTCCAGTG | |
| | | BacCow-257p | **FAM-**TAGGGGTTC**/ZEN/**TGAGAGGAAGGTCCCCC-**IBFQ** | |
| | CowM2 | CowM2F | CGGCCAAATACTCCTGATCGT | (16) |
| | | CowM2R | GCTTGTTGCGTTCCTTGAGATAAT | |
| | | CowM2P | **FAM-**AGGCACCTA**/ZEN/**TGTCCTTTACCTCATCAACTACAGACA**-IBFQ** | |
| | CowM3 | CowM3F | CCTCTAATGGAAAATGGATGGTATCT | (16) |
| | | CowM3R | CCATACTTCGCCTGCTAATACCTT | |
| | | CowM3P | **FAM-**TTATGCATT**/ZEN/**GAGCATCGAGGCC**-IBFQ** | |
| | BacBov1 | BacBov-F1 | AAGGATGAAGGTTCTATGGATTGTAAA | (2) |
| | | BacBov-R1 | GAGTTAGCCGATGCTTATTCATACG | |
| | | BacBov-TP1 | **FAM-**ATACGGGAATAAAACC-**MGB-NFQ** | |
| | BacBov2 | BacBov-F2 | GGATTACAGCCCTACGGGTTTTA | (2) |
| | | BacBov-R2 | GGAGTTAGCCGATGCTTATTCATATAA | |
| | | BacBov-TP2 | **FAM-**AAGAGTAATGTGCACTACGTG-**MGB-NFQ** | |
| Cow mtDNA | Cow-ND5 | Cow-ND5_forward | CAGCAGCCCTACAAGCAATGT | (10) |
| | | Cow-ND5_reverse | GAGGCCAAATTGGGCGGATTAT | |
| | | Cow-ND5_probe | **FAM-**CATCGGCGA**/ZEN/**CATTGGTTTCATTTTAG**-IBFQ** | |

**Table S1.** Continued.

| Target Organism | Assay Name | Primer/Probe Name | Primer sequence (5' --> 3') | Reference |
|---|---|---|---|---|
| Ruminant | | | | |
| Ruminant *Bacteroides* | BacR | BacR_f | GCGTATCCAACCTTCCCG | (17) |
| | | BacR_r | CATCCCCATCCGTTACCG | |
| | | BacR_p | **FAM-**CTTCCGAAAGGGAGATT**-MGB-NFQ** | |
| | Run-2-Bac | BacB2-590F | ACAGCCCGCGATTGATACTGGTAA | (18) |
| | | Bac708Rm | CAATCGGAGTTCTTCGTGAT | |
| | | BacB2-626P | **FAM**-ATGAGGTGG/**ZEN**/ATGGAATTCGTGGTGT-**IBFQ** | |
| Pig | | | | |
| Pig *Bacteroides* | Pig-1-Bac | Pig-1-Bac32Fm | AACGCTAGCTACAGGCTTAAC | (19) |
| | | Pig-1-Bac108R | CGGGCTATTCCTGACTATGGG | |
| | | Pig-1-Bac44P | **FAM**-ATCGAAGCT/**ZEN**/TGCTTTGATAGATGGCG-**IBFQ** | |
| | Pig-2-Bac | Pig-2-Bac41F | GCATGAATTTAGCTTGCTAAATTTGAT | (19) |
| | | Pig-2-Bac163R | ACCTCATACGGTATTAATCCGC | |
| | | Pig-2-Bac113P | **FAM**-TCCACGGGATAGCC-**MGB-NFQ** | |
| Pig mtDNA | Pig-ND5 | Pig-ND5_forward | ACAGCTGCACTACAAGCAATGC | (10) |
| | | Pig-ND5_reverse | GGATGTAGTCCGAATTGAGCTGATTAT | |
| | | Pig-ND5_probe | **FAM-**CATCGGAGA**/ZEN/**CATTGGATTTGTCCTAT**-IBFQ** | |
| Poultry | | | | |
| *Brevibacterium sp.* LA35 | LA35 | LA35F | ACCGGATACGACCATCTGC | (20, 21) |
| | | LA35R | TCCCCAGTGTCAGTCACAGC | |
| | | LA35_Probe | **FAM-**CAGCAGGGA**/ZEN/**AGAAGCCTTCGGGTGACGGTA**-IBFQ** | |

**Table S1. Continued.**

| Target Organism | Assay Name | Primer/Probe Name | Primer sequence (5' --> 3') | Reference |
|---|---|---|---|---|
| Avian | | | | |
| *Lactobacillus spp.* | Av4143 | Av4143F | TGCAAGTCGAACGAGGATTTCT | (22) |
| | | Av4143R | TCACCTTGGTAGGCCGTTACC | |
| | | Av4143P | **FAM-**AGGTGGTTT**/ZEN/**TGCTATCGCTTT**-IBFQ** | |
| Goose | | | | |
| Goose *Bacteroides* | CGOF1-Bac1 | CG1F | GTAGGCCGTGTTTTAAGTCAGC | (23) |
| | | CG1R | AGTTCCGCCTGCCTTGTCTA | |
| | | CG1P | **FAM-**CCGTGCCGT**/ZEN/**TATACTGAGACACTTGAG**-IBFQ** | |
| | CGOF1-Bac2 | CG2F | ACTCAGGGATAGCCTTTCGA | (23) |
| | | CG2R | ACCGATGAATCTTTCTTTGTCTCC | |
| | | CG2P | **FAM-**AATACCTGA**/ZEN/**TGCCTTTGTTTCCCTGCA**-IBFQ** | |
| Goose mtDNA | Goose ND2 | Goose-ND2_forward | CTAACATCCAAATCCCTCGACCCA | (14) |
| | | Goose-ND2_reverse | TCCTATTCAGCCTCCTAGTGCTCT | |
| | | Goose-ND2_probe | **FAM-**TACTCACCG**/ZEN/**CCATAGCCCTAGCCT**-IBFQ** | |
| Gull | | | | |
| *Catellicoccus marimammalium* | Gull2Taqman | Gull2f | TGCATCGACCTAAAGTTTTGAG | (12, 13) |
| | | Gull2r | GTCAAAGAGCGAGCAGTTACTA | |
| | | Gull2p | **FAM-**CTGAGAGGG**/ZEN/**TGATCGGCCACATTGGGACT**-IBFQ** | |
| | Gull-4 | qGull7F | CTTGCATCGACCTAAAGTTTTGAG | (24) |
| | | qGull8R | GGTTCTCTGTATTATGCGGTATTAGCA | |
| | | qGull7P | **FAM-**ACACGTGGG**/ZEN/**TAACCTGCCCATCAGA**-IBFQ** | |
| | LeeSeaGull | CaT#998F | AGGTGCTAATACCGCATAATACAGAG | (25) |
| | | CaT#998R | GCCGTTACCTCACCGTCTA | |
| | | CaT#998P | **FAM-**TTCTCTGTTGAAAGGCGCTT**-MGB-NFQ** | |

**Table S1.** Continued.

| Target Organism | Assay Name | Primer/Probe Name | Primer sequence (5' --> 3') | Reference |
|---|---|---|---|---|
| Deer, Beaver, Muskrat | | | | |
| Deer mtDNA | Deer cytb | Deer-cytb_forward | TAACCCGATTCTTCGCCTTCCTCT | (14) |
| | | Deer-cytb_reverse | GTCTGCGTCTGATGGAATTCCTGAT | |
| | | Deer-cytb_probe | **FAM-**CCTCCCATT**/ZEN/**TATCATCGCAGCACTTGCT**-IBFQ** | |
| Beaver *Bacteroides* | Beapol01 | Beapol-F02 | AGCATTTTTCAAGCTTGCTT | (26) |
| | | Beapol-R01 | ACTTAATGCCATCCCGTATTAA | |
| | | Beapol-P | **FAM-**CAACCTACC**/ZEN/**GTTTACTCTCGG**-IBFQ** | |
| Muskrat *Bacteroides* | MuBa01 | MuBa01F_mod | AACYTCTTTTGTCAGGGAACA | (27) This study |
| | | MuBa01R_mod | GACATTTTACCGCTGACTTGA | |
| | | MuBa01P_mod | **FAM-**TGAGTGTAC**/ZEN/**CTGAAGAAAAAGCATCGGC**-IBFQ** | |

**Table S2.** Primer and probe sequences used in this study to quantify fecal indicator bacteria, pathogenic bacteria, protozoans, DNA virus, and internal amplification controls (DNA targets)

| Target Organism | Assay Name | Primer/Probe Name | Primer sequence (5' --> 3') | Reference |
|---|---|---|---|---|
| Fecal indicator bacteria | | | | |
| *Enterococcus* spp. | Entero1 | ECST748F | GAGAAATTCCAAACGAACTTG | (28) |
| | | ENC854R | CAGTGCTCTACCTCCATCATT | |
| | | GPL813TQ | **FAM-**TGGTTCTCT**/ZEN/**CCGAAATAGCTTTAGGGCTA**-IBFQ** | |
| *E. coli* | *uidA* | uidA405F | CAACGAACTGAACTGGCAGA | (29) |
| | | uidA525R | CATTACGCTGCGATGGAT | |
| | | uidA429P | **FAM-**CCCGCCGGG**/ZEN/**AATGGTGATTAC**-IBFQ** | |
| Bacterial pathogens | | | | |
| Pathogenic *E. coli* | *stx*$_1$ | VS1 (Forward) | CATAGTGGAACCTCACGACGCAGT | (30) |
| | | VS2 (Reverse) | TTTGCCGAAAACGTAAAGCTTCA | |
| | | VS3 (Probe) | **FAM-**TGTGGCAAG**/ZEN/**AGCGATGTTACGGTTTG**-IBFQ** | |
| | *stx*$_2$ | Stx2F | ACCACATCGGTGTCTGTTATTAACC | (31) |
| | | Stx2R | CGGTAGAAAGTATTTGTTGCCGTATT | |
| | | Stx2P | **FAM-**TTTGCTGTG**/ZEN/**GATATACGAGGGCTTGATGTCTAT**-IBFQ** | |
| | *eaeA* | VS8 (Forward) | GGCGGATTAGACTTCGGCTA | (30) |
| | | VS9 (Reverse) | CGTTTTGGCACTATTTGCCC | |
| | | VS10 (Probe) | **FAM-**AACGCCGAT**/ZEN/**ACCATTACTTATACCGCGACG**-IBFQ** | |
| *Shigella* spp. | *ipaH* | ipaH-U1 | CCTTTTCCGCGTTCCTTGA | (32) |
| | | ipaH-L1 | CGGAATCCGGAGGTATTGC | |
| | | ipaH_Probe | **FAM-**CGCCTTTCC**/ZEN/**GATACCGTCTCTGCA**-IBFQ** | |
| | *virA* | Forward | CTGCATTCTGGCAATCTCTT | (33) |
| | | Reverse | GGGACAACTGCGTTGATTTT | |
| | | virA_Probe | **FAM-**TGCCCAAAT**/ZEN/**TATGTCCGAAAACA**-IBFQ** | |

**Table S2.** Continued

| Target Organism | Assay Name | Primer/Probe Name | Primer sequence (5' --> 3') | Reference |
|---|---|---|---|---|
| *Salmonella* spp. | *invA* | invA_176F | CAACGTTTCCTGCGGTACTGT | (34) |
| | | invA_291R | CCCGAACGTGGCGATAATT | |
| | | invA_Tx_208 | **FAM-**CTCTTTCGT/**ZEN**/CTGGCATTATCGATCAGTACCA-**IBFQ** | |
| | *ttr* | ttr-6_forward | CTCACCAGGAGATTACAACATGG | (35) |
| | | ttr-4_reverse | AGCTCAGACCAAAAGTGACCATC | |
| | | ttr-5 | **FAM-**CACCGACGG**/ZEN/**CGAGACCGACTTT**-IBFQ** | |
| *Campylobacter jejuni* | *ciaB* | ciaB_Forward | CAACTTTATATTTGCACTCCGATG | (36) |
| | | ciaB_Reverse | GGAACGACTTGAGCTGAGAATAAAC | |
| | | ciaB_Probe | **FAM-**TATGGTGCTGAACTTAAA**-MGB-NFQ** | |
| | *hipO* | CJ_hipO_F | AATGCACAAATTTGCCTTATAAAAGC | (37) |
| | | CJ_hipO_R | TNCCATTAAAATTCTGACTTGCTAAATA | |
| | | CJ_hipO_probe | **FAM-**ACATACTACTTCTTTATTGCTTG**-MGB-NFQ** | |
| *Campylobacter coli* | *cadF* | CC_cadF_F | GAGAAATTTTATTTTTATGGTTTAGCTGGT | (37) |
| | | CC_cadF_R | ACCTGCTCCATAATGGCCAA | |
| | | CC_cadF_probe | **FAM-**CCTCCACTT**/ZEN/**TTATTATCAAAAGCGCCTTTAGAAA**-IBFQ** | |
| *Clostridium perfringens* | *cpe* | cpe F | AACTATAGGAGAACAAAATACAATAG | (38) |
| | | cpe R | TGCATAAACCTTATAATATACATATTC | |
| | | cpe Pr | **FAM-**TCTGTATCT**/ZEN/**ACAACTGCTGGTCCA**-IBFQ** | |
| | CPerf16S | CPerf165F | CGCATAACGTTGAAAGATGG | (39) |
| | | CPerf269R | CCTTGGTAGGCCGTTACC | |
| | | CPerf187FAM | **FAM**-TCATCATTC/ZEN/AACCAAAGGAGCAATCC-**IBFQ** | |
| *Listeria monocytogenes* | *iap* | iap (forward) | CTCAAATACGAATGCTAACCAAGGT | (40) |
| | | iap (reverse) | TTTGAGCTTCAGCAATAATAGCACTT | |
| | | iap (probe) | **FAM-**TAACAGCAATTCAAGCGC**-MGB-NFQ** | |

**Table S2.** Continued.

| Target Organism | Assay Name | Primer/Probe Name | Primer sequence (5' --> 3') | Reference |
|---|---|---|---|---|
| *Listeria monocytogenes* | *hlyA* | hlyA-LisM-F | ACTGAAGCAAAGGATGCATCTG | (41) |
| | | hlyA-LisM-R | TTTTCGATTGGCGTCTTAGGA | |
| | | hlyA-LisM-P | **FAM-**CACCACCAG**/ZEN/**CATCTCCGCCTGC**-IBFQ** | |
| *Vibrio cholerae* | *ctxA* | VC_ctxAF | TTTGTTAGGCACGATGATGGAT | (42) |
| | | VC_ctxAR | ACCAGACAATATAGTTTGACCCACTAAG | |
| | | VC_ctxA_probe | **FAM-**TGTTTCCAC**/ZEN/**CTCAATTAGTTTGAGAAGTGCCC**-IBFQ** | |
| | *toxR* | VC_toxR_420/334F | GTTTGGCGWGAGCAAGGTTT | (43, 44) |
| | | VC_toxR_585R | TCTCTTCTTCAACCGTTTCCA | |
| | | toxR_464/378FAM | **FAM**-CGCAGAGTMGAAATGGCTTGG-**MGB-NFQ** | |
| *Vibrio parahaemolyticus* | *tdhS* | VP_tdhF | AAACATCTGCTTTTGAGCTTCCA | (45) |
| | | VP_tdhR | CTCGAACAACAAACAATATCTCATCAG | |
| | | VP_tdhS | **FAM**-TGTCCCTTT/**ZEN**/TCCTGCCCCCGG-**IBFQ** | |
| *Legionella* spp. | *ssrA* | PanLeg-F | GGCGACCTGGCTTC | (46) |
| | | PanLeg-R1 | GGTCATCGTTTGCATTTATATTTA | |
| | | PanLeg-P1 | **FAM-**CGTGGGTTGCAA**-MGB-NFQ** | |
| *L. pneumophila* | *mip* | Lp-F | TTGTCTTATAGCATTGGTGCCG | (46) |
| | | Lp-R | CCAATTGAGCGCCACTCATAG | |
| | | Lp-P | **FAM-**CGGAAGCAA**/ZEN/**TGGCTAAAGGCATGCA**-IBFQ** | |
| *L. pneumophila* serogroup 1 | *wzm* | Lp1-F | TGCCTCTGGCTTTGCAGTTA | (46) |
| | | Lp1-R | CACACAGGCACAGCAGAAACA | |
| | | Lp1-P | **FAM-**TTTATTACTCCACTCCAGCGAT**-MGB-NFQ** | |
| *Mycobacterium* spp. | *atpE* | FatpE | CGGYGCCGGTATCGGYGA | (47) |
| | | RatpE | CGAAGACGAACARSGCCAT | |
| | | PatpE | **FAM-**ACSGTGATG**/ZEN/**AAGAACGGBGTRAA**-IBFQ** | |

**Table S2.** Continued.

| Target Organism | Assay Name | Primer/Probe Name | Primer sequence (5' --> 3') | Reference |
|---|---|---|---|---|
| *M. avium* complex | MAC ITS | MAC_Forward | TTGGGCCCTGAGACAACACT | (48) |
| | | MAC_Reverse | GCAACCACTATCCAATACTCAAACAC | |
| | | MAC_Probe | **FAM-**CCGTGTGGA**/ZEN/**GTCCCTCCATCTTGG**-IBFQ** | |
| Protozoan pathogens | | | | |
| *Giardia* sp. | beta-Giardin P241 | P241_Forward | CATCCGCGAGGAGGTCAA | (49) |
| | | P241_Reverse | GCAGCCATGGTGTCGATCT | |
| | | P241_Probe | **FAM-**AAGTCCGCC**/ZEN/**GACAACATGTACCTAACGA**-IBFQ** | |
| *Cryptosporidium* spp. | COWP P702 | P702_Forward | CAAATTGATACCGTTTGTCCTTCTG | (49) |
| | | P702_Reverse | GGCATGTCGATTCTAATTCAGCT | |
| | | P702_Probe | **FAM-**TGCCATACA**/ZEN/**TTGTTGTCCTGACAAATTGAAT**-IBFQ** | |
| *Entamoeba histolytica* | Ehf | Ehf | AACAGTAATAGTTTCTTTGGTTAGTAAAA | (50) |
| | | Ehr | CTTAGAATGTCATTTCTCAATTCAT | |
| | | Ehp | **FAM-**ATTAGTACA**/ZEN/**AAATGGCCAATTCATTCA**-IBFQ** | |
| *Acanthamoeba spp.* | Acant | AcantF900 | CCCAGATCGTTTACCGTGAA | (51) |
| | | AcantR1100 | TAAATATTAATGCCCCCAACTATCC | |
| | | AcantP1000 | **FAM-**CTGCCACCG**/ZEN/**AATACATTAGCATGG**-IBFQ** | |
| *Naegleria fowleri* | NaegI | NaeglF192 | GTGCTGAAACCTAGCTATTGTAACTCAGT | (51) |
| | | NaeglR344 | CACTAGAAAAAGCAAACCTGAAAGG | |
| | | NfowlP | **FAM-**ATAGCAATA**/ZEN/**TATTCAGGGGAGCTGGGC**-IBFQ** | |
| Viral pathogen (DNA virus) | | | | |
| Human Adenovirus | ADV | JTVX(F) | GGACGCCTCGGAGTACCTGAG | (52) |
| | | JTVX(R) | ACIGTGGGGTTTCTGAACTTGTT | |
| | | JTVX(P) | **FAM-**CTGGTGCAG/**ZEN**/TTCGCCCGTGCCA-**IBFQ** | |

**Table S2.** Continued.

| Target Organism | Assay Name | Primer/Probe Name | Primer sequence (5' --> 3') | Reference |
|---|---|---|---|---|
| Internal amplification control | | | | |
| Internal amplification control | HF183-IAC | HF183 | ATCATGAGTTCACATGTCCG | (3) |
| | | BacR287 | CTTCCTCTCAGAACCCCTATCC | |
| | | BacP234IAC | **FAM-**AACACGCCGTTGCTACA**-MGB-NFQ** | |
| *Pseudogulbenkiania* sp. strain NH8B-1D2 | NH8B_3960tnp | NH8B_3960tnp_1841F | CTGGCTGTCTAGGCCCTGTCT | (53) |
| | | NH8B_3960tnp_1901R | CCTGCAGGCATGCAAGCT | |
| | | NH8B_3960tnp_1863 | **FAM-**TTATACACATCTCAACCCTG**-MGB-NFQ** | |

**Table S3.** Primer and probe sequences used in this study to quantify RNA targets.

| Target Organism | Assay Name | Primer/Probe Name | Primer/Probe Sequence (5' --> 3') | Reference |
|---|---|---|---|---|
| Viral pathogens (RNA viruses) | | | | |
| Aichivirus | AiV Total | AiV-AB-F | GTCTCCACHGACACYAAYTGGAC | (54) |
| | | AiV-AB-R | GTTGTACATRGCAGCCCAGG | |
| | | AiV-AB-TP | **FAM-**TTYTCCTTYGTGCGTGC**-MGB-NFQ** | |
| Astrovirus | AsV | AV1 | CCGAGTAGGATCGAGGGT | (55) |
| | | AV2 | GCTTCTGATTAAATCAATTTTAA | |
| | | AVs | **FAM-**CTTTTCTGT**/ZEN/**CTCTGTTTAGATTATTTTAATCACC**-IBFQ** | |
| Enteroviruses | EV | EV2 | CCCCTGAATGCGGCTAATC | (56) |
| | | EV1 | GATTGTCACCATAAGCAGC | |
| | | EV-probe | **FAM-**CGGAACCGA**/ZEN/**CTACTTTGGGTGTCCGT**-IBFQ** | |
| Human Norovirus | HNoV-GI | NIFG1F | ATGTTCCGCTGGATGCG | (57) |
| | | NV1LCR | CCTTAGCCATCATCATTTAC | |
| | | NIFGIP | **FAM-**TGTGGACAG**/ZEN/**GAGAYCGCRATCT**-IBFQ** | |
| | HNoV-GII | NIFG2F | ATGTTCAGRTGGATGAGRTTCTC | (57) |
| | | COG2R | TCGACGCCATCTTCATTCACA | |
| | | QNIFS | **FAM-**AGCACGTGG**/ZEN/**GAGGGCGATCG**-IBFQ** | |
| | HNoV-GIV | NIFG4F | ATGTACAAGTGGATGCGRTTC | (57) |
| | | COG2R | TCGACGCCATCTTCATTCACA | |
| | | NIFG4P | **FAM-**AGCACTTGG**/ZEN/**GAGGGGGATCG**-IBFQ** | |
| Rotavirus A | RoV | JVKF | CAGTGGTTGATGCTCAAGATGGA | (58) |
| | | JVKR | TCATTGTAATCATATTGAATACCCA | |
| | | JVKP | **FAM-**ACAACTGCA**/ZEN/**GCTTCAAAAGAAGWGT**-IBFQ** | |

**Table S3.** Continued.

| Target Organism | Assay Name | Primer/Probe Name | Primer/Probe Sequence (5' --> 3') | Reference |
|---|---|---|---|---|
| Sapovirus GI, GII, GIV, and GV | SaV | HuSaV-F1 | GGCHCTYGCCACCTAYAAYG | (59) |
| | | HuSaV-F2 | GACCARGCHCTCGCYACCTAYGA | |
| | | HuSaV-F3 | GCWRYKGCWTGYTAYAACAGC | |
| | | HuSaV-R | CCYTCCATYTCAAACACTA | |
| | | HuSaV-TP-a | **FAM-**CCNCCWATRWACCA**-MGB-NFQ** | |
| | | HuSaV-TP-b | **FAM-**CCNCCWACRWACCA**-MGB-NFQ** | |
| Hepatitis A virus | HAV | HAV68 | TCACCGCCGTTTGCCTAG | (60) |
| | | HAV241 | GGAGAGCCCTGGAAGAAAG | |
| | | HAV150(-) | **FAM-**CCTGAACCTGCAGGAATTAA**-MGB-NFQ** | |
| Hepatitis E virus | HEV | HECOM-S | CGGCGGTGGTTTCTGGRGTG | (61) |
| | | HECOM-AS | GGGCGCTKGGMYTGRTCNCGCCAAGNGGA | |
| | | TP-HECOM | **FAM-**CCCCYATAT/**ZEN**/TCATCCAACCAACCCCTTYGC**-IBFQ** | |
| Human MST marker (RNA viruses) | | | | |
| Pepper Mild Mottle Virus | PMMV | PMMV-FP-rev | GAGTGGTTTGACCTTAACGTTTGA | (62, 63) |
| | | PMMV-RP | TTGTCGGTTGCAATGCAAGT | |
| | | PMMV-Probe | **FAM-**CCTACCGAAGCAAATG**-MGB-NFQ** | |
| Process control (RNA viruses) | | | | |
| Murine norovirus | MNV | MNV-S | CCGCAGGAACGCTCAGCAG | (64) |
| | | MNV-AS | GGYTGAATGGGGACGGCCTG | |
| | | MNV-TP | **FAM-**ATGAGTGATGGCGCA**-MGB-NFQ** | |

**Table S4.** gBlock sequences used as DNA standards. Primer and probe annealing sites are shown in red and blue, respectively.

| Assay Name | gBlock Sequence | Genbank Accession |
|---|---|---|
| MST markers | | |
| AllBac / BacGeneral | TAGGGGTT**CTGAGAGGAAGGTCCCCCAC**ATTGGAACTGAGACACGGTCCAAACTCCTACGGGAGGC**AGCAGTGAGGAATATT**GGTCAATGGGCGCTAGC**CTGAACCAGCCAAGTAGCGTG**AAGGATGA | NR_074784 |
| HF183 | CCAGGATGGG**ATCATGAGTTCACATGTCCG**CATGATTAAAGGTATTTTCCGGTAGACGATG**GGGATGCGTTCCATTAG**ATAGTAGGCGGGGTAACGGCCCACCTAGTCAACGAT**GGATAGGGGTTCTGAGAGGAAG**GTCCCCCACATTGGA | AB242142 |
| gyrB | TCCTATGTCAGGT**GGCGGTCTTCCGGGTAAA**C**TGGCCGACTGCTC**GGACAAAGACCCGCAGAAGTGTGAGTTATTCCTCGTCGAGGGAGACTCTGCCGGCGGTACAGCTAAGCAAGGTCGTAACCGTGCATTTCAGGCTATTCTTCCACTACGCGGTAAGATTCTGAACGTAGAGAAAGCCATGTATCACAAAGCGCTTGAAAGCGAAGAAATACGCAATATATACACGGCACTGGGTGTCACTATCGGAACGGAAGAAGACAGCAAAGCTGCCAATATTGATAAGCTGCGCTATCATAAAATCATTATCATGACCGATGCCGACGTCGATGGATCACACATCGACACACTGATCATGA**CTTTTTTCTTCCGCTATATGCCA**CAGATCATCCAGAATG | AB017713 |
| HumM2 | CCGCAAAGTC**CGTCAGGTTTGTTTCGGTATTG**AG**TATCGAAAATCTCACGGATTAACTCTTGTGTACGC**TCCGGACGACCG**GCTAACGCATATAAATAAGTTACGTGATGA**CTCGGTTCGT | CP022412 |
| H8 | GCTTGGCCTG**ACAGTCAGCGAGATTCTTC**GCCACGCCGGCGTGGCGCATCTGCTGCTGGAGGCGGACGCGCAGAAGGTCGAGGCCGCGCGTGCCGCCGGCGCGCCG**GTGTTCCATGCCGATGCCAGT**CGGCCCGATACCTTGCTGGCTGCCGGCTTGACGCATGCACAC**TTGGTGGTGCTGACGTTC**GCCCATGCCC | KY887595 |
| H12 | GGAAATACGTC**GTAAAAGGACTGCCGGGAAA**GGCGTTGTTTACCATATTAGCGCAGCCACAATGTCGCTTGATGATGAGAGAAACAG**GGCTAAGGGCGAAGCGTTTACTCT**CAACGACAGTGATCGCCAGCTAAGCCTCTGGGTGCAGCTGTATCGCAATATGTCGCCAGCTTCGCGGGCGCAACTGATGGAGCAGGCCCTCAGG**CTGGTAAAGGACGATCTGA**TGTCTCAGGG | CP056396 |
| Mnif | GATATTTTATGTGTT**GAAAGCGGAGGTCCTGAA**CCGGGTGTTGGCTGTG**CCGGACGTGGTGTAACAGTAGCTA**TGAAAAGACTTGAAAACTTAGGTGTTTTTGATAAGGATTTGGATGTAGTCATTTATGGTGTACTTGGAGATGTT**GTTTGCGGAGGTTTTTCAGT**GCCTTTACGTGAAAAGTATG | AB019138 |

**Table S4.** Continued.

| Assay Name | gBlock Sequence | Genbank Accession |
|---|---|---|
| CPQ_056 | TGCTGAACAAACTGCTAATG**CAGAAGTACAAACTCCTAAAAAACGTAGAG**GTAGAGGTATT**AATAACGATTTACGTGATGTAAC**TCGTAAAAAGTTTGATGAACGTACTGATTGTAATAAA**GCTAATGGCTTGTTTATTGGTCATC**TTGAAGATGTTAAAGTTG | NC_024711 |
| CPQ_064 | GATATTGGTGGGGAGAT**TGTATAGATGCTGCTGCAACTGTACTC**T**CTGAAATTGTTCATAAGCAA**ATTGATATTTCTATTAAAAGTCAATTTCTATTTGTTCTTAAACATATTGCTTATACTTTTAGAAATATTATTT**ATGGACAAGATAAAGATGAAAACAACG**ATTATAATATTGCTAGG | JQ995537 |
| Human-ND5 | GATGCCAACA**CAGCAGCCATTCAAGCAATCC**TATACAACCG**TATCGGCGATATCGGTTTCATCCTCG**CCTTAGCATGATTTATCCTACACTCCAACTCATGAGACCCACAACAAATAGCCCTTCTAAACGCTAATCCAAGCCTCACCCCACTACTAGGCCTCCTCCTAGCAGCAGCAGG**CAAATCAGCCCAATTAGGTCTCCACC**CCTGACTCCC | AY972053 |
| BacCan-UCD / DogBact | TACGTGTAGG**CGCTTGTATGTACCGGTACG**AATAAGCATCGGCTAACTCCGTGCCAGCAGCCGCGGTAATACGGAGGATGCGAGCGTTATCCGGATTTATTGGGTTTAAAG**GGAGCGCAGACGGGTTTT**TAAGTCAGCTGTGAAAGTTTGGGGCTCAACCTTAAAATTGCAGTTGATNCTGGAGACCTTGAGTGCAGTTGAGGCAGGCGGA**ATTCGTGGTGTAGCGGTGAAATGCTTAGATATCACGAAGAACTCCGATTG** | AY695700 |
| Dog-ND5 | CGCATTAACA**GGCATGCCTTTCCTTACAGGATTC**TACTCCAAAGACCTGATCATCGAGTCCGCTAACACGTCGAATACCAACGCCTGAGCCCT**CTTAATTACACTCGTTGCCACATCCC**TAACCGCTGC | EU078704 |
| BacCow-UCD | AACGCGTAT**CCAACCTTCCCGTTACTC**TGGGATAGCCTTCCGAAAGGGAGATTAATACCGGATGGTGTTCAAGTGGTGCATATTATTTGAACTAAAGATTTATCGGTAACGGATGGGGATGCGTACCATTAGATAGTAGGCGGGGTAACGGCCCACCTAGTCCACGATGGT**TAGGGGTTCTGAGAGGAAGGTCCCCC**A**CACTGGAACTGAGACACGGTCC**AGACTCCTAC | NR_112936 |
| CowM2 | ATCG**CGGCCAAATACTCCTGATCGT**ACTCGAGAT**AGGCACCTATGTCCTTTACCTCATCAACTACAGACA**AA**ATTATCTCAAGGAACGCAACAAGC**ATCG | |
| CowM3 | ATCG**CCTCTAATGGAAAATGGATGGTATCT**TTGGAGCCTTTGAAAGCACTCGAGCC**TTATGCATTGAGCATCGAGGCC**GGAAAGCAGGAACTTATATATAAT**AAGGTATTAGCAGGCGAAGTATGG**ATCG | |
| BacBov1 | AAGTAGCGTG**AAGGATGAAGGTTCTATGGATTGTAAA**CTTCTTTT**ATACGGGAATAAAACC**TCCCACGTGTGGGAGCTTGTATGTAC**CGTATGAATAAGCATCGGCTAACTC**CGTGCCAGCA | AY597142 |
| BacBov2 | GTAGCGTGAA**GGATTACAGCCCTACGGGTTTTA**AACTTCTTTTATAT**AAGAGTAATGTGCACTACGTGT**AGTGTATTGCAAGTA**TTATATGAATAAGCATCGGCTAACTCC**GTGCCAGCAG | |
| Cow-ND5 | GATGCAAACA**CAGCAGCCCTACAAGCAATCT**TATATAACCG**CATCGGCGACATTGGTTTCATTTTAG**CAATAGCATGGTTCCTAACAAATCTCAATACCTGAGACCTCCAACAGATCTTCATACTAAACCCAAGCGACTCAAACATACCCTTGATTGGACTAGCATTAGCTGCAACCGG**AAAATCCGCCCAATTTGGCCTC**CACCCGTGAC | NC_006853 |

**Table S4.** Continued.

| Assay Name | gBlock Sequence | Genbank Accession |
|---|---|---|
| BacR | GGTGCGTAACGCGTATCCAACCTTCCCGTTACTCAGGGATAGCCTTCCGAAAGGGAGATTAAT ACCTGATGGTATTCAAATTTCGCATGTTATTTGAACTAAAGATTTATCGGTAACGGATGGGGATG CGTGACATTA | AF233400 |
| Run-2-Bac | TAGCTCAACCACAGCCCGCGATTGATACTGGTAACCTTGAGTGCGGATGAGGTGGATGGAATT CGTGGTGTAGCGGTGAAATGCTTAGATATCACGAAGAACTCCGATTG | EU913520 |
| Pig-1-Bac | TGGCTCAGGATGAACGCTAGCTACAGGCTTAACACATGCAAGTCGAGGGGCAGCATTATCGAA GCTTGCTTTGATAGATGGCGACCGGCGCACGGGTGAGTAACGCGTATCCAACCTTCCCCATAG TCAGGAATAGCCCGGCGAAAGTCGAATTAATG | EU797142 |
| Pig-2-Bac | GTCGCGGGGCAGCATGAATTTAGCTTGCTAAATTTGATGGCGACCGGCGCACGGGTGAGTAAC GCGTATCCAACCTTCCCCTGTCCACGGGATAGCCCGTCGAAAGGCGGATTAATACCGTATGAG GTCACACGACGGCATCAGAGTG | EU797137 |
| Pig-ND5 | AGACGCCAACACAGCTGCACTACAAGCAATCCTATACAACCGCATCGGAGACATTGGATTTGT CCTATCCATAGCATGATTCCTAACCCACTCAAACGCATGAGATCTTCAACAAATCTTTATACTAAA CAATGAATGCCCAAACATACCATTAATCGGCCTACTCCTAGCTGCAGCAGGAAAATCAGCTCAA TTCGGACTACATCCCTGATTGCCC | AF034253 |
| LA35 | GAAACTGGGTCTAATACCGGATACGACCATCTGCCGCATGGCGGGTGGTGGAAAGTTTTTCGA TTGGGGATGGGCTCGCGGCCTATCAGTTTGTTGGTGGGGTAATGGCCTACCAAGGCGACGACG GGTAGCCGGCCTGAGAGGGCGACCGGCCACACTGGGACTGAGACACGGCCCAGACTCCTACG GGAGGCAGCAGTGGGGAATATTGCACAATGGGGGAAACCCTGATGCAGCGACGCAGCGTGCG GGATGACGGCCTTCGGGTTGTAAACCGCTTTCAGCAGGGAAGAAGCCTTCGGGTGACGGTAC CTGCAGAAGAAGTACCGGCTAACTACGTGCCAGCAGCCGCGGTAATACGTAGGGTACGAGCGT TGTCCGGAATTATTGGGCGTAAAGAGCTCGTAGGTGGTTGGTCACGTCTGCTGTGGAAACGCAA CGCTTAACGTTGCGCGGGCAGTGGGTACGGGCTGACTAGAGTGCAGTAGGGGAGTCTGGAATT CCTGGTGTAGCGGTGAAATGCGCAGATATCAGGAGGAACACCGGTGGCGAAGGCGGGACTCT GGGCTGTGACTGACACTGGGGAGCGAAAGCATGGGG | FJ462358 |
| Av4143 | GGCGGCGTGCCTAATACATGCAAGTCGAACGAGGATTTCTTACACTGAGTGCTTGCACTCACC GTAAGAAATTCGAGTGGCGGACGGGTGAGTAACACGTGGGTAACCTGCCCAAAAGAAGGGGAT AACATTTGGAAACAAATGCTAATACCGTATAACCATGATGACCGCATGGTCATTATGTAAAAGGT GGTTTTGCTATCGCTTTTGGATGGACCCGCGGCGTATTAACTAGTTGGTAGGGTAACGGCCTAC CAAGGTGATGATACGTAGCCGAGTT | LN864462 |

**Table S4.** Continued.

| Assay Name | gBlock Sequence | Genbank Accession |
|---|---|---|
| CGOF1-Bac1 | TAAAGGGTGC**GTAGGCCGTGTTTTAAGTCAGC**GGTGAAAAGTTGCAGCTCAACTGTAA**CCGTGCCGTTATACTGAGACACTTGAG**TG**TAGACAAGGCAGGCGGAACT**CGTAGTGTAG | GU222166 |
| CGOF1-Bac2 | CCTTCCCTTT**ACTCAGGGATAGCCTTTCGA**AAGAAAGATT**AATACCTGATGCCTTTGTTTCCCTGCA**TGG**GGAGACAAAGAAAGATTCATCGGT**AAAGGATGGG | GU222167 |
| Goose ND2 | AACCCTACTCCTA**CTAACATCCAAATCCCTCGACCCA**GCCC**TACTCACCGCCATAGCCCTAGCC**TCA**ACAGCACTAGGAGGCTGAATAGGA**TTAAACCAAACACAAACACG | NC_007011 |
| Gull2Taqman | AACTGATGCT**TGCATCGACCTAAAGTTTTGAG**TGGCGGACGGGTGAGTAACACGTGGGTAACCTGCCCATCAGAGGGGGACAACACTTGGAAACAGGTGCTAATACCGCATAATACAGAGAACCGCATGGTTCTTTGTTGAAAGGCGCTTCTGGTGTCGCTGATGGATGGACCCGCGGTGCATTAGCTAGACGGTGAGGTAACGGCTCACCGTGGCAATGATGCATAGCCGAC**CTGAGAGGGTGATCGGCCACATTGGGACT**GAGACACGGCCCAAACTCCTACGGGAGGCAGCAGTAGGGAATCTTCGGCAATGGACGAAAGTCTGACCGAGCAACGCCGCGTGAGTGAAGAAGGTTTTCGGATCGTAAAACTCTGTTGTTAGAGAAGAACAGGAGCGA**TAGTAACTGCTCGCTCTTTGAC**GGTATCTAACCAG | NR_042357 |
| Gull-4 | CTGATGAACTGATG**CTTGCATCGACCTAAAGTTTTGAG**TGGCGGACGGGTGAGTA**ACACGTGGGTAACCTGCCCATCAGA**GGGGGACAACACTTGGAAACAGG**TGCTAATACCGCATAATACAGAGAACC**GCATGGTTCTTTGTTG | NR_042357 |
| LeeSeaGull | GACAACACTTGGAAAC**AGGTGCTAATACCGCATAATACAGAG**AACCGCATGG**TTCTTTGTTGAAAGGCGCTT**CTGGTGTCGCTGATGGATGGACCCGCGGTGCATTAGC**TAGACGGTGAGGTAACGGC**TCACCGTGGCAATGATGCATAG | NR_042357 |
| Deer cytb | AAAGCAACCC**TAACCCGATTCTTCGCCTTCCACT**TTAT**CCTCCCATTTATCATCGCAGCACTTGCT**ATAGTCCATTTACTCTTCCTCCACGAAACAGGATCTAACAACCC**AACAGGAATTCCATCAGACGCAGAC**AAAATTCCATTC | DQ379370 |
| Beapol01 | GTCGAGGGGC**AGCATTTTTCAAGCTTGCTT**GAAAAGATGGCGACCGGCGCACGGGTGAGTAACACGTATC**CAACCTACCGTTTACTCTCGG**ATAGCCTTCCGAAAGGAAGA**TTAATACGGGATGGCATTAAGT**TTTCACATGG | KC243716 |
| MuBa01 | TAGTATTGTA**AACTTCTTTTGTCAGGGAACA**AAAAAGGTTATGTATAACCCTC**TGAGTGTACCTGAAGAAAAAGCATCGGGC**TAACTCCGTGCCAGCAGCCGCGGTAATACGGAGGATGCGAGCGTTATCCGGATTTATTGGGTTTAAAGGGTGCGCAGGCGGCAAG**TCAAGTCAGCGGTAAAATGTC**GGGGCTCAAC | JF703174 |

**Table S4.** Continued.

| Assay Name | gBlock Sequence | Genbank Accession |
|---|---|---|
| Fecal indicator bacteria | | |
| Entero1 | GTGGGTAGCG**GAGAAATTCCAAACGAACTTG**GAGATAGC**TGGTTCTCTCCGAAATAGCTTTAGGGCTA**GCCTCGGAATTGAG**AATGATGGAGGTAGAGCACTG**TTTGGACTAG | NR_076202 |
| *uidA* | TTTGTGTGAA**CAACGAACTGAACTGGCAGA**CTAT**CCCGCCGGGAATGGTGATTAC**CGACGAAAACGGCAAGAAAAAGCAGTCTTACTTCCATGATTTCTTTAACTATGCCGGA**ATCCATCGCAGCGTAATG**CTCTACACCA | S69414 |
| Bacterial pathogens | | |
| *stx1* | TTATCTGGATTTAATGTCG**CATAGTGGAACCTCACTGACGCAGT**C**TGTGGCAAGAGCGATGTTACGGTTTG**TTACTGTGACAGC**TGAAGCTTTACGTTTTCGGCAAA**TACAGAGGGGATTTCGTACAACACTGGATGAT | M16625 |
| *stx2* | TTGAACATATATCTCAGGGG**ACCACATCGGTGTCTGTTATTAACC**ACACCCCACCGGGCAGTTAT**TTTGCTGTGGATATACGAGGGCTTGATGTCTAT**CAGGCGCGTTTTGACCATCTTCGTCTGATTATTGAGCAAAATAATTTATATGTGGCCGGGTTCGTT**AATACGGCAACAAATACTTTCTACCG**TTTTTCAGATTTTACACAT | X07865 |
| *eaeA* | TCAAGTTGTCGACCAGGTTGGGGTAACGGACTTTAC**GGCGGATAAGACTTCGGCTA**AAGCGGAT**AACGCCGATACCATTACTTATACCGCGACG**GTGAAAAAGAATGGGGTAGCTCAGGCTAATGTCCCTGTTTCATTTAATATTGTTTCAGGAACTGCAACTCTTG**GGGCAAATAGTGCCAAAACG**GATGCTAACGGTAAGG | X60439 |
| *ipaH* | GAACATGAAGAGCATGCCAACA**CCTTTTCCGCGTTCCTTGA**C**CGCCTTTCCGATACCGTCTCTGCA**C**GCAATACCTCCGGATTCCG**TGAACAGGTCGCTGCATGGCTGGAAAAACTCAGTGCCTCTGCGGAGCT | M32063 |
| *virA* | GACCCTGCATTTATATCTGCACTAACAT**CTGCATTCTGGCAATCTCTT**CACATCACGTCTTCCTCTGTGGAACACATATAT**TGCCCAAATTATGTCCGAAAACA**TAGAAAACAGGCTTAATTTCATGCCTGAACAACGAGTTATTAACAATTGTGGACATATTATA**AAAATCAACGCAGTTGTCCC**TAAAAACGACACTGCAATCTCTGCCTCTG | AF047364 |
| *invA* | CTCAGTTTTT**CAACGTTTCCTGCGGTACTGT**TAATTACCACG**CTCTTTCGTCTGGCATTATCGATCAGTACCA**GTCGTCTTATCTTGATTGAAGCCGATGCCGGTGA**AATTATCGCCACGTTCGGG**CAATTCGTTA | NC_003197 |
| *ttr* | GCGCTACTGATTATTATTCGTGAAACGCTGAACGGA**CTCACCAGGAGATTACAACATGG**CTAATTTAACCCGTCGTCAGTGGCTA**AAAGTCGGTCTCGCCGTCGGTG**G**GATGGTCACTTTTGGTCTGAGCT**ACCGTGATGTGGCGAAACGCGCAATTG | AF282268 |

**Table S4.** Continued.

| Assay Name | gBlock Sequence | Genbank Accession |
|---|---|---|
| *ciaB* | CTCATCATTTGGAACGACTTGAGCTGAGAATAAACCTTTAAGTTCAGCACCATAAAATATCATCGGAGTGCAAATATAAAGTTGAGTTTTTTCT | CP000025 |
| *hipO* | AGATTTAAAGTGCCATTAAAATTCTGACTTGCTAAATACTTTGCAGCAAGCAATAAAGAAGTAGTATGTCCATCATGACCGCAAGCATGCATTACATTTTCTTTTTTGCTTTTATAAGGCAAATTTGTGCATTCTTGCAAAGG | CP000025 |
| *cadF* | CATTGATTTAGGTGAGAAATTTTATTTTTATGGTTTAGCTGGTGGGGGATATGAGGATTTTTCTAAAGGCGCTTTTGATAATAAAAGTGGAGGATTTGGCCATTATGGAGCAGGTTTAAAATTTCGCCTT | FJ946041 |
| *cpe* | TGTAGAATATGGATTTGGAATAACTATAGGAGAACAAAATACAATAGAAAGATCTGTATCTACAACTGCTGGTCCAAATGAATATGTATATTATAAGGTTTATGCAACTTATAGAAAGTATCAAG | X81849 |
| CPerf16S | AGATTAATACCGCATAACGTTGAAAGATGGCATCATCATTCAACCAAAGGAGCAATCCGCTATGAGATGGACCCGCGGCGCATTAGCTAGTTGGTGGGGTAACGGCCTACCAAGGCGACGATGCG | NR_121697 |
| *iap* | GAATAAGCTTTTCCAAGGTGTTTTTGAGCTTCAGCAATAATAGCACTTGCACTTGAATTGCTGTTATTGTTGGAAGAACCTTGATTAGCATTCGTATTTGAGTTTGTATTCGTATTAGTATTTGAGTTTGTATTAG | AL591824 |
| *hlyA* | ACTTATCGATTTCATCCGCGTGTTTCTTTTCGATTGGCGTCTTAGGACTTGCAGGCGGAGATGCTGGTGGTGCCATGGATGAAATTAAATTTTCTTTATTGAATGCAGATGCATCCTTTGCTTCAGTTTGTTGCGCAATTG | CP045749 |
| *ctxA* | TCAGACGGGATTTGTTAGGCACGATGATGGATATGTTTCCACCTCAATTAGTTTGAGAAGTGCCCACTTAGTGGGTCAAACTATATTGTCTGGTCATTCTACTT | NC_002505 |
| *toxR* | GCATGACTTTGTTTGGCGAGAGCAAGGTTTTGAAGTCGATGATTCCAGCTTAACCCAAGCCATTTCGACTCTGCGCAAAATGCTCAAAGATTCGACAAAGTCCCCACAATACGTCAAAACGGTTCCGAAACGCGGTTACCAATTGATCGCCCGAGTGGAAACGGTTGAAGAAGAGATGGCTCGCGA | NC_002505 |
| *tdhS* | GCTGCATTCAAAACATCTGCTTTTGAGCTTCCATCTGTCCCTTTTCCTGCCCCCGGTTCTGATGAGATATTGTTTGTTGTTCGAGATACAACTTT | NC_004605 |
| *ssrA* | CAGAATGGGGGGCGACCTGGCTTCGACGTGGGTTGCAAAACCGGAAGTGCATGCCGAGAAGGAGATCTCTCGTAAATAAGACTCAATTAAATATAAATGCAAACGATGAAAACTTTGCTGG | AE017354 |
| *mip* | CAAGGATAAGTTGTCTTATAGCATTGGTGCCGATTTGGGGAAGAATTTTAAAAATCAAGGCATAGATGTTAATCCGGAAGCAATGGCTAAAGGCATGCAAGACGCTATGAGTGGCGCTCAATTGGCTTTAACCGA | AJ810179 |
| *wzm* | TGGTTTAGCCCACACAGGCACAGCAGAAACAGGGTAAATCGCTGGAGTGGAGTAATAAAATAACTGCAAAGCCAGAGGCATAAAATGTTT | AM778127 |

**Table S4.** Continued.

| Assay Name | gBlock Sequence | Genbank Accession |
|---|---|---|
| atpE | GTGGCGCCAT**CGGCGCCGGTATCGGTGA**CGGTGTCGCCGGTAACGCGCTTATCTCCGGTGTCGC<br>CCGGCAACCCGAGGCGCAAGGGCGGCTG**TTCACACCGTTCTTCATCACCGT**CGGTTTGGTTGAGG<br>CGGCATACTTCATCAACCTGGCGTTT**ATGGCGCTGTTCGTCTTCG**CTACACCCGT | AL123456 |
| MAC ITS | AGACACACTA**TTGGGCCCTGAGACAACACT**CGGTCCGT**CCGTGTGGAGTCCCTCCATCTTGG**TGG<br>TGGGGTGTG**GTGTTTGAGTATTGGATAGTGGTTGC**GAGCATCTAG | CP000479 |
| Protozoan pathogens | | |
| beta-Giardin P241 | GCCAGGCCTCGTTCGAGGA**CATCCGCGAGGAGGTCAA**G**AAGTCCGCCGACAACATGTACCTAAC**<br>**GA**TCAAGGAGG**AGATCGACACCATGGCTGC**AAACTTCCGCAAGTCCCTTGCGGAGATGGGCGACA<br>CACTCAACAACGTTGAG | M36728 |
| COWP P702 | GAATGGCACATGTAAATTAATTCAA**CAAATTGATACCGTTTGTCCTTCTG**GTTTTGTTGAAGAAGGAA<br>ATAGATGTGTTCAATATCTCCCTGCAAATAAAATCTGTCCTCCTGG**ATTCAATTTGTCAGGACAACA**<br>**ATGTATGGCA**CCAGAATC**AGCTGAATTAGAATCGACATGCC**CACCTAATTCAATATTTGAAAATGG | AF248743 |
| Ehf | GGCTCATTAT**AACAGTAATAGTTTCTTTGGTTAGTAAAA**TACAAGGATAGCTTTGTGAATGATAAAG<br>ATAATACTTGAGACGATCCAGTTTGT**ATTAGTACAAAATGGCCAATTCATTCAATGAATTGAGAAAT**<br>**GACATTCTAAG**TGAGTTAGGATGCCACG | KT253454 |
| Acant | GTCCCTGGGG**CCCAGATCGTTTACCGTGAA**AAAATTAGAGTGTTCAAAGCAGGCAGATCCAATTTT<br>**CTGCCACCGAATACATTAGCATGG**GATAATGGAATAGGACCCTGTCCTCCTATTTTCAGTTGGTTTT<br>GGCAGCGCGAGGACTAGGGTAATGATTAATAG**GGATAGTTGGGGGCATTAATATTTA**ATTGTCAGA<br>G | U07416 |
| Naegl | AGTAGTATTT**GTGCTGAAACCTAGCTATTGTAACTCAGT**TTCTCTGGGT**ATAGCAATATATTCAGGG**<br>**GAGCTGGGC**ATCGACCGCTAGCAGGTGCCCGCAAGGGCGCGGGAAAGTGAGCTAACAAGGTTTTC<br>ATAAGG**CCTTTCAGGTTTGCTTTTTCTAGTG**GCCAGGCAGA | U80059 |
| Viral pathogen (DNA virus) | | |
| ADV | TCGCCGGGCA**GGACGCCTCGGAGTACCTGAG**CCCGGGC**CTGGTGCAGTTCGCCCGTGCCA**CCGA<br>TACCTACTTCAGCCTGGGG**AACAAGTTCAGAAACCCCACCGT**GGCTCCCACC | X51782 |
| Internal amplification control | | |
| HF183-IAC | CCAGGATGGG**ATCATGAGTTCACATGTCCG**CATGATTAAAGGTATTTTCCGGTAGACGATG**TGTAG**<br>**CAACGGCGTGTT**ATAGTAGGCGGGGTAACGGCCCACCTAGTCAACGAT**GGATAGGGGTTCTGAGA**<br>**GGAAG**GTCCCCCACA | Ref. (3) |
| NH8B_3960tnp | AGCCAACATC**CTGGCTGTCTAGGCCCTGTCT**C**TTATACACATCTCAACCCTGA**A**AGCTTGCATGCCT**<br>**GCAGG**TCGACTCTAG | Ref. (53) |

**Table S4.** Continued.

| Assay Name | gBlock Sequence | Genbank Accession |
|---|---|---|
| Viral pathogens (RNA viruses) | | |
| AiV Total | TATCTTCCCTGTCTCCACTGACACCAATTGGACCTCCACCTCCAGTCCTACCGCATATCCCCTCCCTTTCTCCTTCGTGCGTGCTTACCCAGACTCCTCCTGGGCTGCCATGTACAACACCCACTCCA | AB040749 |
| AsV | CTGGAGACCGCGGCCACGCCGAGTAGGATCGAGGGTACAGTCTCCTTTACTTTTCTGTCTCTGTTTAGATTATTTTAATCACCATTTAAAATTGATTTAATCAGAAGCAAAAA | MN433705 |
| EV | AATCCTCCGGCCCCTGAATGCGGCTAATCCCAACCTCGGGGCAGGTGGTCACAAACCAGTGATTGGCCTGTCGTAACGCGCAAGTCCGTGGCGGAACCGACTACTTTGGGTGTCCGTGTTTCCTTTTATTTTATTGTGGCTGCTTATGGTGACAATCACAGATTGTT | MW080377 |
| HNoV-GI | ATGGCAGGCCATGTTCCGCTGGATGCGCTTCCATGACCTCGGATTGTGGACAGGAGATCGCGATCTTCTGCCCGAATTCGTAAATGATGATGGCGTCTAAGGACGCTACATC | M87661 |
| HNoV-GII | ACAAGAGCCAATGTTCAGATGGATGAGATTCTCAGATCTGAGCACGTGGGAGGGCGATCGCAATCTGGCTCCCAGCTTTGTGAATGAAGATGGCGTCGAATGACGCCAA | X86557 |
| HNoV-GIV | TGTGCCAAAGTTTGAGTCTATGTACAAGTGGATGCGATTCTCAGACCTGAGCACTTGGGAGGGGGATCGCGATCTCGCTCCCGATTTTGTGAATGAAGATGGCGTCGAGTGACGCTGC | AF414427 |
| RoV | GGCTTTTAATGCTTTTCAGTGGTTGATGCTCAAGATGGAGTCTACTCAACAGATGGCATCTTCTATTATTAACTCTTCTTTTGAAGCTGCAGTTGTCGCTGCAACTTCTACATTAGAATTAATGGGTATTCAATATGATTATAATGAAGTATACACTAGAG | DQ146697 |
| SaV | GAACACCAATTATGATCACGCTCTCGCCACCTACAACGCATGGTTCATAGGTGGTACAGTGCCTGACCCAGAACGCCCCACTGAAGGCGCGTCCAAAATAGTGTTTGAGATGGAGGGCAATGGCTCCCAACAAGGGGCACGACCAAAAAGTCCACCTCAAAGTGTTGACCTTCCTGGCACGGTTGGCCCG | DQ366345 |
| HAV | ACTTGATACCTCACCGCCGTTTGCCTAGGCTATAGGCTAAATTTTCCCTTTCCCTTTTCCCTTTCCTATTCCCTTTGTTTTGCTTGTAAATATTAATTCCTGCAGGTTCAGGGTTCTTAAATCTGTTTCTCTATAAGAACACTCATTTTTCACGCTTTCTGTCTTCTTTCTTCCAGGGCTCTCCCCTTGCCCTA | M14707 |
| HEV | GCGGCGGTTCCGGCGGTGGTTTCTGGGGTGACCGGGTTGATTCTCAGCCCTTCGCAATCCCCTATATTCATCCAACCAACCCCTTCGCCCCCGATGTCACCGCTGCGGCCGGGGCTGGACCTCGTGTTCGCCAACCCGCCCGACCACTCGGCTCCGCTTGGCGTGACCAGGCCCAGCGCCCTCACGTCGTA | M73218 |
| Human MST marker (RNA viruses) | | |
| PMMoV | ATTAGGCGTAGATCCATTGGTGGCAGCAAAGGTAATGGTAGCTGTGGTTTCAAATGAGAGTGGTTTGACCTTAACGTTTGAGAGGCCTACCGAAGCAAATGTCGCACTTGCATTGCAACCGACAATTACATCAAAGGAGGAAGG | NC_003630 |

**Table S4.** Continued.

| Assay Name | gBlock Sequence | Genbank Accession |
|---|---|---|
| Process control (RNA viruses) | | |
| MNV | TGCTGAGACC**CCGCAGGAACGCTCAGCAG**TCTTTGTGAATGAGG**ATGAGTGATGGCGCA**GCGCCAAAAGCCAACGGCTCTGAAGCCAGCGGCCAGGATCTTGTTCCTACCGCCGTTGAA**CAGGCCGTCCCCATTCAGCC**CGTGGCTGGC | AB435514 |

**Table S5.** Amplification efficiency (E), limit of quantification (LOQ), and r$^2$ value of the MFQPCR assays.

| | Target Organism | Assay | Efficiency (%) | LOQ (copies/μL) | r² |
|---|---|---|---|---|---|
| MST marker | *Bacteroides* spp. | All Bac | 93.0 | 20 | 0.998 |
| | | BacGeneral | 90.7 | 20 | 0.999 |
| | Human | HF183 | 96.0 | 200 | 0.995 |
| | | gyrB | 71.0 | 2 | 0.984 |
| | | HumM2 | 94.8 | 2 | 0.995 |
| | | H8 | 79.1 | 2 | 0.989 |
| | | H12 | 92.6 | 2 | 0.982 |
| | | Mnif | 100.7 | 2 | 0.989 |
| | | CPQ_056 | 90.3 | 2 | 0.995 |
| | | CPQ_064 | 92.1 | 2 | 0.991 |
| | | Human-ND5 | 83.5 | 2 | 0.986 |
| | | PMMoV | 100.2 | 2 | 0.995 |
| | Dog | BacCan-UCD | 93.9 | 2 | 0.995 |
| | | DogBact | 86.0 | 2 | 0.981 |
| | | Dog-ND5 | 97.9 | 2 | 0.995 |
| | Cow | CowM2 | 99.4 | 2 | 0.996 |
| | | CowM3 | 95.8 | 2 | 0.993 |
| | | BacBov1 | 89.9 | 2 | 0.998 |
| | | BacBov2 | 89.7 | 2 | 0.998 |
| | | Cow-ND5 | 86.5 | 2 | 0.993 |
| | Ruminant | BacR | 96.5 | 2 | 0.997 |
| | | Run-2-Bac | 88.6 | 2 | 0.997 |
| | Pig | Pig-1-Bac | 91.4 | 2 | 0.990 |
| | | Pig-2-Bac | 92.1 | 2 | 0.994 |
| | | Pig-ND5 | 91.1 | 2 | 0.996 |
| | Avian | Av4143 | 80.1 | 2 | 0.988 |
| | Goose | CGOF1-Bac1 | 89.2 | 2 | 1.000 |
| | | CGOF1-Bac2 | 101.6 | 2 | 0.990 |
| | | Goose ND2 | 100.2 | 2 | 0.992 |
| | Gull | Gull-4 | 86.3 | 2 | 0.993 |
| | | LeeSeaGull | 102.4 | 2 | 0.996 |
| | Deer | Deer cytb | 95.1 | 2 | 0.995 |
| | Beaver | Beapol01 | 80.8 | 2 | 0.987 |
| | Muskrat | MuBa01 | 85.1 | 2 | 0.991 |

**Table S5.** Continued.

| | Target Organism | Assay | Efficiency (%) | LOQ (copies/µL) | $r^2$ |
|---|---|---|---|---|---|
| FIB | *Enterococcus* spp. | Entero1 | 88.5 | 2 | 0.989 |
| | *Escherichia coli* | uidA | 87.1 | 2 | 0.996 |
| Bacterial pathogen | Pathogenic *E. coli* | stx1 | 86.8 | 2 | 0.994 |
| | | stx2 | 94.2 | 2 | 0.994 |
| | | eaeA | 104.7 | 2 | 0.988 |
| | *Shigella* spp. | ipaH | 99.8 | 2 | 0.995 |
| | | virA | 93.2 | 2 | 0.994 |
| | *Salmonella* spp. | invA | 102.9 | 2 | 0.994 |
| | | ttr | 90.0 | 20 | 0.994 |
| | *Campylobacter jejuni* | ciaB | 102.4 | 2 | 0.996 |
| | | hipO | 93.7 | 2 | 0.997 |
| | *Campylobacter coli* | cadF | 104.9 | 2 | 0.990 |
| | *Clostridium perfringens* | CPerf16S | 98.4 | 2 | 0.989 |
| | | cpe | 93.5 | 2 | 0.986 |
| | *Listeria monocytogenes* | iap | 98.8 | 2 | 0.995 |
| | | hlyA | 103.7 | 2 | 0.991 |
| | *Vibrio cholerae* | ctxA | 93.1 | 20 | 0.995 |
| | | toxR | 102.0 | 2 | 0.981 |
| | *Vibrio parahaemolyticus* | tdhS | 86.8 | 2 | 0.974 |
| | *Legionella* spp. | ssrA | 101.8 | 2 | 0.993 |
| | *Le. pneumophila* | mip | 95.2 | 2 | 0.998 |
| | *Le. pneumophila* ser. 1 | wzm | 97.8 | 2 | 0.997 |
| | *Mycobacterium* spp. | atpE | 80.8 | 20 | 0.989 |
| | *M. avium* complex | MAC ITS | 93.2 | 2 | 0.998 |
| Protozoan pathogen | *Giardia* spp. | beta-Giardin p241 | 105.8 | 2 | 0.989 |
| | *Cryptosporidium* spp. | COWP P702 | 92.4 | 2 | 0.994 |
| | *Entamoeba histolytica* | Ehf | 92.7 | 2 | 0.993 |
| | *Acanthamoeba* spp. | Acant | 97.5 | 2 | 0.988 |
| | *Naegleria fowleri* | NaegI | 96.0 | 2 | 0.994 |

**Table S5.** Continued.

| | Target Organism | Assay | Efficiency (%) | LOQ (copies/µL) | $r^2$ |
|---|---|---|---|---|---|
| Viral pathogen | Human Adenovirus | ADV | 114.1 | 2000 | 0.998 |
| | Aichivirus | AiV | 104.3 | 2 | 0.985 |
| | Astrovirus | AsV | 84.6 | 2 | 0.995 |
| | Enteroviruses | EV | 95.2 | 2 | 0.988 |
| | Human norovirus | HNoV-GI | 102.4 | 2 | 0.984 |
| | | HNoV-GIV | 100.7 | 2 | 0.988 |
| | Rotavirus A | RoV | 85.1 | 20 | 0.985 |
| | Sapovirus | SaV | 84.4 | 2 | 0.981 |
| | Hepatitis A virus | HAV | 91.6 | 2 | 0.988 |
| | Hepatitis E virus | HEV | 77.0 | 2 | 0.991 |
| Control | Internal amplification control | HF183-IAC | 125.2 | 200 | 0.992 |
| | *Pseudogulbenkiania* sp. NH8B-2D9 | NH8B_3960tnp | 105.0 | 2 | 0.995 |
| | Murine norovirus | MNV | 97.9 | 2 | 0.993 |

**Table S6.** Statistical significance (p value) of Kendall's rank correlations between gene copy numbers and water quality parameters. Values smaller than 0.05 are highlighted in red.

| | AllBac | atpE | BacBov1 | Bac General | cpe | CPerf16S | CPQ_056 | CPQ_064 | Entero1 | Gull_4 | gyrB | H8 | Human_ ND5 | HumM2 | LeeSea Gull | Mnif | PMMoV | ssrA | uidA | pH | CONDUCT | TURBID | TOT_ COLIFORM | E_COLI | WATER_ TEMP |
|---|---|---|---|---|---|---|---|---|---|---|---|---|---|---|---|---|---|---|---|---|---|---|---|---|---|
| AllBac | 0.000 | | | | | | | | | | | | | | | | | | | | | | | | |
| atpE | 0.110 | 0.000 | | | | | | | | | | | | | | | | | | | | | | | |
| BacBov1 | 0.707 | 0.290 | 0.000 | | | | | | | | | | | | | | | | | | | | | | |
| BacGeneral | **0.000** | 0.138 | 0.779 | 0.000 | | | | | | | | | | | | | | | | | | | | | |
| cpe | 0.290 | 0.448 | **0.018** | 0.338 | 0.000 | | | | | | | | | | | | | | | | | | | | |
| CPerf16S | **0.012** | 0.110 | 0.245 | **0.018** | 0.138 | 0.000 | | | | | | | | | | | | | | | | | | | |
| CPQ_056 | 0.290 | 0.169 | 0.110 | 0.245 | 0.391 | **0.026** | 0.000 | | | | | | | | | | | | | | | | | | |
| CPQ_064 | 0.338 | 0.205 | 0.138 | 0.290 | 0.338 | **0.037** | **0.000** | 0.000 | | | | | | | | | | | | | | | | | |
| Entero1 | **0.018** | 0.138 | 0.391 | **0.026** | 0.110 | **0.002** | **0.037** | **0.026** | 0.000 | | | | | | | | | | | | | | | | |
| Gull_4 | 0.707 | 0.926 | 0.391 | 0.779 | 0.852 | 0.852 | 0.448 | 0.508 | 0.638 | 0.000 | | | | | | | | | | | | | | | |
| gyrB | 0.290 | 0.852 | 0.707 | 0.245 | 0.926 | 0.290 | 0.638 | 0.572 | 0.448 | 0.852 | 0.000 | | | | | | | | | | | | | | |
| H8 | 0.391 | 0.245 | **0.007** | 0.338 | 0.138 | 0.138 | 0.050 | 0.067 | 0.448 | 0.110 | 0.926 | 0.000 | | | | | | | | | | | | | |
| Human_ND5 | 0.205 | 0.067 | 0.338 | 0.245 | 0.508 | 0.205 | 0.205 | 0.245 | 0.338 | 0.448 | 0.779 | 0.205 | 0.000 | | | | | | | | | | | | |
| HumM2 | 0.087 | 0.067 | 0.338 | 0.169 | 0.205 | 0.138 | 0.290 | 0.338 | 0.067 | 0.707 | 0.926 | 0.391 | 0.391 | 0.000 | | | | | | | | | | | |
| LeeSeaGull | 0.638 | 0.852 | 0.572 | 0.572 | 0.779 | 0.926 | 0.638 | 0.707 | 0.707 | 0.110 | 0.638 | 0.508 | 0.138 | 0.926 | 0.000 | | | | | | | | | | |
| Mnif | 0.110 | 0.087 | 0.391 | 0.138 | 0.338 | **0.007** | **0.018** | **0.012** | **0.004** | 0.926 | 0.448 | 0.245 | 0.169 | 0.245 | 1.000 | 0.000 | | | | | | | | | |
| PMMoV | 0.448 | 0.138 | 0.391 | 0.391 | 0.338 | 0.448 | 0.067 | 0.050 | 0.138 | 0.779 | 1.000 | 0.169 | 0.110 | 0.169 | 0.707 | 0.087 | 0.000 | | | | | | | | |
| ssrA | 1.000 | 0.779 | **0.026** | 0.926 | 0.110 | 0.338 | 0.110 | 0.087 | 0.508 | 0.290 | 0.572 | 0.067 | 0.572 | 0.852 | 0.572 | 0.391 | 0.391 | 0.000 | | | | | | | |
| uidA | 0.050 | 0.067 | 0.338 | **0.037** | 0.205 | 0.050 | 0.087 | 0.110 | **0.018** | 0.852 | 0.926 | 0.138 | 0.205 | **0.012** | 0.779 | 0.067 | 0.110 | 0.852 | 0.000 | | | | | | |
| pH | 0.779 | 0.338 | 0.448 | 0.572 | 0.779 | 0.779 | 0.779 | 0.852 | 1.000 | 0.707 | 0.050 | 0.638 | 0.508 | 0.638 | 0.508 | 0.852 | 0.448 | 0.852 | 0.926 | 0.000 | | | | | |
| CONDUCT | 0.169 | 0.290 | 0.779 | 0.205 | 0.338 | 0.338 | 0.707 | 0.779 | 0.205 | 0.926 | 0.707 | 0.572 | 0.572 | 0.245 | 0.572 | 0.290 | 0.290 | 0.508 | 0.110 | 0.448 | 0.000 | | | | |
| TURBID | 0.448 | 0.138 | 0.779 | 0.391 | 0.707 | 0.707 | 0.852 | 0.926 | 0.638 | 0.926 | 0.572 | 0.852 | 0.338 | 0.448 | 1.000 | 0.638 | 0.779 | 0.638 | 0.338 | 0.245 | 0.205 | 0.000 | | | |
| TOT_COLIFORM | 0.245 | 0.638 | 0.290 | 0.205 | **0.037** | 0.169 | 0.572 | 0.508 | 0.290 | 0.638 | 0.448 | 0.338 | 0.707 | 0.448 | 0.707 | 0.391 | 0.779 | 0.508 | 0.245 | 0.852 | 0.138 | 0.508 | 0.000 | | |
| E_COLI | 0.852 | 0.638 | 0.391 | 0.926 | 0.245 | 0.572 | 0.338 | 0.290 | 0.290 | 0.638 | 1.000 | 0.572 | 0.448 | 0.448 | 0.852 | 0.205 | 0.138 | 0.779 | 0.245 | 0.448 | 0.205 | 0.926 | 0.205 | 0.000 | |
| WATER_TEMP | 0.474 | 0.741 | 0.183 | 0.536 | 0.095 | 0.149 | 0.415 | 0.360 | 0.360 | 0.360 | 0.474 | 0.310 | 0.962 | 0.814 | 0.670 | 0.360 | 0.670 | 0.120 | 0.536 | 0.474 | 0.670 | 0.814 | 0.056 | 0.741 | 0.000 |
